# Prediction Errors Disrupt Hippocampal Representations and Update Episodic Memories

**DOI:** 10.1101/2020.09.29.319418

**Authors:** Alyssa H. Sinclair, Grace M. Manalili, Iva K. Brunec, R. Alison Adcock, Morgan D. Barense

## Abstract

The brain supports adaptive behavior by generating predictions, learning from errors, and updating memories to incorporate new information. *Prediction error,* or surprise, triggers learning when reality contradicts expectations. Prior studies have shown that the hippocampus signals prediction errors, but the hypothesized link to memory updating has not been demonstrated. In a human fMRI study, we elicited mnemonic prediction errors by interrupting familiar narrative videos immediately before the expected endings. We found that prediction error reversed the relationship between univariate hippocampal activation and memory: greater hippocampal activation predicted memory preservation after expected endings, but memory updating after surprising endings. In contrast to previous studies, we showed that univariate activation was insufficient for understanding hippocampal prediction error signals. We explained this surprising finding by tracking both the evolution of hippocampal activation patterns and connectivity between the hippocampus and neuromodulatory regions. We found that hippocampal activation patterns stabilized as each narrative episode unfolded, suggesting sustained episodic representations. Prediction errors disrupted these sustained representations, and the degree of disruption predicted memory updating. The relationship between hippocampal activation and subsequent memory depended on concurrent basal forebrain activation, supporting the idea that cholinergic modulation regulates attention and memory. We conclude that prediction errors create conditions that favor memory updating, prompting the hippocampus to abandon ongoing predictions and make memories malleable.

**Significance:** Our brains draw on memories to predict the future; when our predictions are incorrect, we must update our memories to improve future predictions. Past studies have demonstrated that the hippocampus signals *prediction error*, or surprise, but have not linked this neural signal to memory updating. Here, we uncover this missing connection: We show that mnemonic prediction errors change the role of the hippocampus, reversing the relationship between hippocampal activation and memory outcomes. We examine the mechanisms of this shift in neural processing, showing that prediction errors disrupt the temporal continuity of hippocampal patterns. We propose that prediction errors disrupt sustained representations and enable memory updating. Our findings bear implications for improving education, understanding eyewitness memory distortion, and treating pathological memories.

## Introduction

In daily life, we continuously draw on past experiences to predict the future. Expectation and surprise shape learning across many situations, such as when we discover misinformation in the news, receive feedback on an exam, or make decisions based on past outcomes. When our predictions are incorrect, we must update our mnemonic models of the world to support adaptive behavior. *Prediction error* is a measure of the discrepancy between expectation and reality; this surprise signal is both evident in brain activity and related to learning (1–6). The brain dynamically reconstructs memories during recall, recreating and revising past experiences based on current information (7). The intuitive idea that surprise governs learning has long shaped our understanding of memory, reward learning, perception, action, and social behavior (2, 8–14). Yet, the neural mechanisms that allow prediction error to update memories remain unknown.

Past research has implicated the hippocampus in each of the mnemonic functions required for learning from prediction errors: retrieving memories to make predictions, identifying discrepancies between past and present, and encoding new information (2, 15–20). Functional MRI (fMRI) studies have shown that hippocampal activation increases after predictions are violated; this surprise response has been termed *mismatch detection* (18, 19, 21–23), or *mnemonic prediction error* (20). These past studies have shown that the hippocampus *detects* mnemonic prediction errors. Several theoretical frameworks have hypothesized that this hippocampal prediction error signal could update memories (17, 20, 24–27), but this crucial link for understanding how we learn from error has not yet been demonstrated.

What mechanisms could link hippocampal prediction errors to memory updating? A leading hypothesis is that prediction errors shift the focus of attention and adjust cognitive processing (20, 28–32). After episodes that align with expectations, we should continue generating predictions and shift attention *internally*, sustaining and reinforcing existing memories. However, after mnemonic prediction errors, we should reset our expectations and shift attention *externally*, preparing to encode new information and update memories. Consistent with this idea, mnemonic prediction errors have been shown to enhance the hippocampal *input* pathway that supports encoding, but suppress the *output* pathway that supports retrieval (20). We propose that surprising events may also change intrinsic hippocampal processing, changing the effect of hippocampal activation on memory outcomes.

Neuromodulation may be a critical factor that regulates hippocampal processing and enables memory updating. Currently, there is mixed evidence supporting two hypotheses: acetylcholine and/or dopamine could act upon the hippocampus to regulate processing after surprising events (24–27, 29, 31, 33, 34). Several models have proposed that acetylcholine from the medial septum (within the basal forebrain) regulates the balance between input and output pathways in the hippocampus (27–29, 35–38), thus allowing stored memories to be compared with perceptual input (31, 38, 39). After prediction errors, acetylcholine release could change hippocampal processing and enhance encoding or memory updating (26, 29, 33, 37, 39). On the other hand, dopamine released from the ventral tegmental area (VTA), if transmitted to the hippocampus, could also modulate hippocampal plasticity after prediction errors. Past studies have shown that the hippocampus and VTA are co-activated after surprising events (40, 41). Other work has shown that co-activation of the hippocampus and VTA predicts memory encoding and integration (42–45). Overall, basal forebrain and VTA neuromodulation are both candidate mechanisms for regulating hippocampal processing and memory updating.

In the present study, we used an fMRI task with human participants to examine trial-wise hippocampal responses to prediction errors during narrative videos. During the **Encoding phase**, participants viewed 70 full-length videos that featured narrative episodes with salient endings (e.g., a baseball batter hitting a home run) (Figure 1A). During the **Reactivation phase** the following day, participants watched the videos again (Figure 1B). We elicited mnemonic prediction errors by interrupting half of the videos immediately before the expected narrative ending (e.g., the video ends while the baseball batter is mid-swing). These surprising interruptions were comparable to the prediction errors employed in prior studies of memory updating (1). Half of the videos were presented in Full-length form (as previously seen during the Encoding phase), and half were presented in Interrupted form (eliciting prediction error).

**Figure 1.**
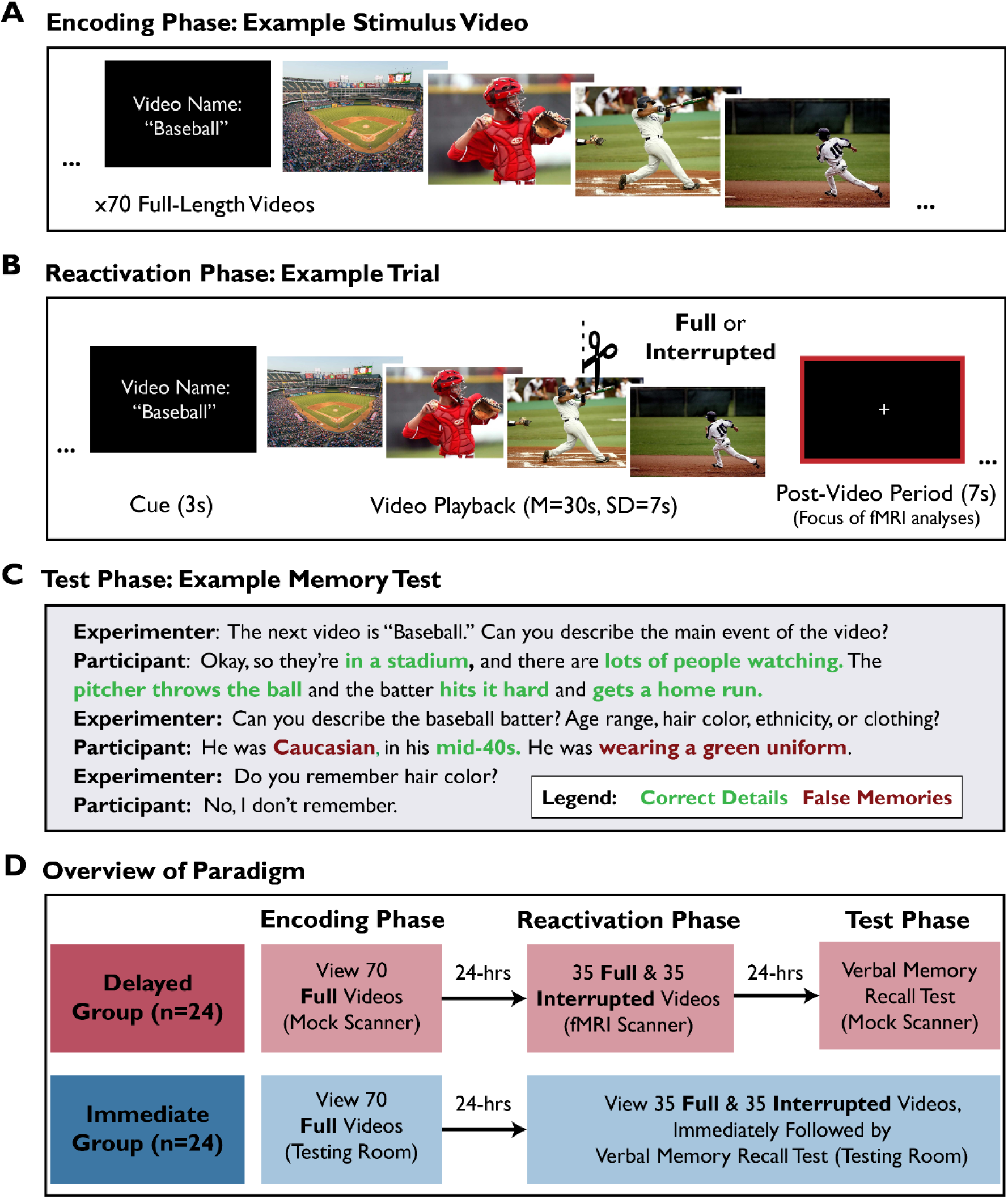
Overview of experimental paradigm. A) During the Encoding phase, all videos were presented in Full-length form. Here we show example frames depicting a stimulus video. B) During the Reactivation phase, participants viewed the 70 videos again, but half (35) were interrupted to elicit mnemonic prediction error. Participants were cued with the video name, watched the video (Full or Interrupted), and then viewed a fixation screen. The “Baseball” video was interrupted when the batter was mid-swing. fMRI analyses focused on the Post-Video fixation periods after each video (red highlighted box). Thus, visual and auditory stimulation were matched across Full and Interrupted conditions, allowing us to compare Post-Video neural activation while controlling for perceptual input. C) During the Test phase, participants answered structured interview questions about all 70 videos, and were instructed to answer based on their memory of the Full video originally shown during the Encoding phase. Here we show example text illustrating the memory test format and scoring of correct details (our measure of memory preservation) and false memories (our measure of memory updating, because false memories indicate that the memory has been modified). The void response (“I don’t remember”) is not counted as a false memory. D) Overview of the experiment. All participants completed Encoding, Reactivation, and Test phases of the study. The Delayed group (fMRI participants) completed the Test phase 24 hours after Reactivation, because prior studies have shown that memory updating becomes evident only after a delay (e.g., to permit protein synthesis). The Immediate group completed the Test phase immediately after Reactivation and was not scanned. The purpose of the Immediate group was to test the behavioral prediction that memory updating required a delay.

During the **Test phase**, participants completed a memory test in the form of a structured interview (Figure 1C). On each trial, participants were cued with the name of the video and recalled the narrative. The experimenter then probed for further details with pre-determined questions (e.g., “Can you describe the baseball batter’s ethnicity, age range, or clothing?”). Our critical measure of memory updating was *false memories*, because the presence of a false memory indicates that the original memory was changed in some way. Although it can be adaptive to update real-world memories by incorporating relevant new information, we expected that our laboratory paradigm would induce false memories because participants would integrate interfering details across similar episodes (1, 7). Because we were interested in false memories as a measure of memory updating, we instructed participants not to guess and permitted them to skip details they could not recall.

Prior research in human and animals has shown that some memory updating effects only emerge after delays that allow protein synthesis to occur during consolidation and reconsolidation (1, 46–48). Therefore, to test our primary question about the neural correlates of memory updating, fMRI participants completed the Encoding, Reactivation, and Test phases over three days, with 24-hours between each session (*Delayed* group, n = 24). In addition, we tested the behavioral prediction that memory updating would *require* a delay (i.e., because transforming a memory trace requires protein synthesis) by recruiting a separate group of participants who completed the Test phase immediately after the Reactivation phase on Day 2 (*Immediate* group, n = 24) (Figure 1D). Delayed group participants completed the Reactivation phase while undergoing an fMRI scan, whereas Immediate group participants (n = 24) were not scanned. Our primary fMRI analyses examined the fixation period immediately following the offset of Full and Interrupted videos (Post-Video period; Figure 1B, right) during the Reactivation phase in the Delayed group. Importantly, this design compares neural responses to surprising and expected video endings while controlling for visual and auditory input.

Our approach allowed us to test several questions set up by the prior literature. First, we used naturalistic video stimuli to examine the effect of mnemonic prediction error on hippocampal activation and episodic memories. Second, to investigate hippocampal processing, we used multivariate analyses to track how episodic representations were sustained or disrupted over time. Third, to test hypotheses about neuromodulatory mechanisms, we related hippocampal activation and memory updating to activation in the basal forebrain and VTA.

## Results

### Behavioral Results

We transcribed and scored memory tests for two key measures: number of unique *correct details* (Figure 2A) and *false memories* (Figure 2B) – reflecting memory preservation and updating, respectively. We also collected *confidence ratings* and scored the number of *forgotten videos* (Supplementary Information, *Confidence and Forgetting*, Figure S1). We defined false memories as distorted details that the participant recalled from the episode (e.g., “The pitcher wore a green uniform”). Void responses (e.g., “I don’t remember”) were not counted as false memories, but were missed opportunities to earn points for correct details. Importantly, our measures for correct details and false memories were independent; there was no limit to the number of details a participant could recall about a video, and each detail was scored as correct or false. We conducted linear mixed-effects regression to predict memory outcomes (either *correct details* or *false memories*) from the factors *group* (Delayed vs. Immediate) and *reactivation type* (Full vs. Interrupted). In all models, we included random effects to account for by-subject and by-video variability (Supplementary Information, *Supplementary Methods, Linear Mixed Effects Regression*).

**Figure 2.**
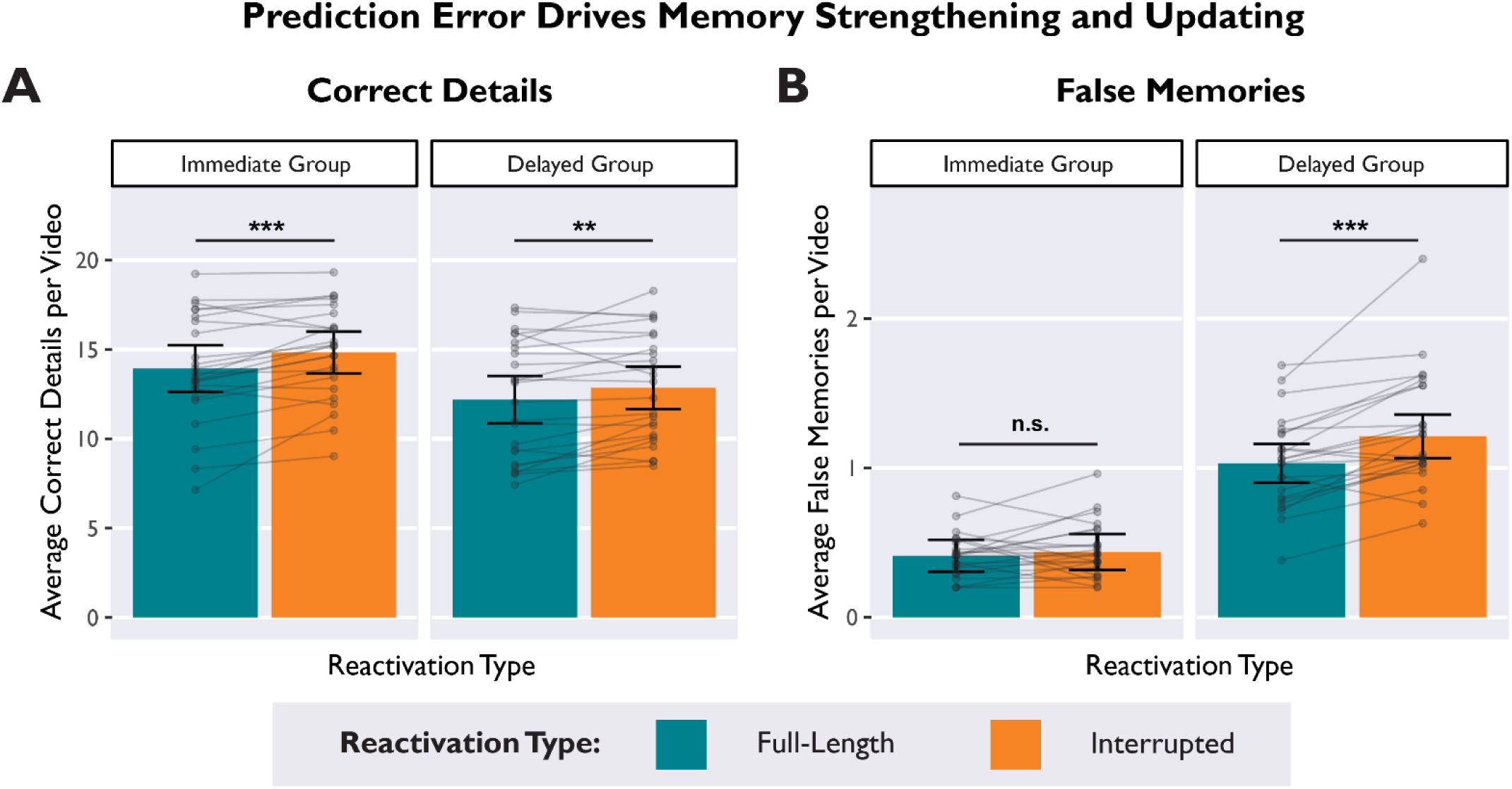
Prediction errors strengthened and updated memories over distinct time-courses. A) In both the Delayed and Immediate groups, average Correct Details were higher for videos that were Interrupted during memory reactivation, demonstrating that prediction error can strengthen memory recall both immediately and after a delay. B) In the Delayed group (but not the Immediate group), average False Memories were higher for videos that were Interrupted during memory reactivation. This interaction demonstrates that prediction error enabled memory updating, but only after a delay. Bars depict estimated marginal means from a linear mixed effects model. Subject averages are overlaid on top to display the distribution: Dots indicate average scores by-participant, and lines connect within-subjects measures. Error bars depict 95% confidence intervals. * *p* < .05, ** *p* < .01, *** *p* < .001.

#### Prediction Error Increased Correct Details

We found that prediction errors during memory reactivation enhanced recall of correct details (main effect of reactivation type: β = -0.07, 95% CI [-0.12, -0.02], *t* = -2.75, *p* = .008) (Figure 2A), such that participants in both groups reported more correct details for Interrupted videos than Full videos (Delayed Group: β = -0.05, *t* = -2.07, *p* = .042; Immediate Group: β = - 0.08, *t* = -2.50, *p* = .015) (Table S1A). Even though the video endings were omitted, prediction errors strengthened and preserved existing memories. Participants in the Delayed group recalled fewer correct details than participants in the Immediate group (main effect of group: β = 0.16, 95% CI [0.01, 0.31], *t* = 2.16, *p* < .036), likely because the Delayed group completed the memory test after 24 hours. There was no interaction between group and reactivation type (β = - 0.01, 95% CI [-0.04, 0.02], *t* = -0.68, *p* = .495), indicating that the effect of prediction error enhancing correct details did not require a delay.

#### Prediction Error Increased False Memories

We found that prediction errors selectively increased false memories in the Delayed group, replicating our past behavioral results (48) (significant interaction between reactivation type and group, β = 0.04, 95% CI [0.01, 0.07], *t* = 2.61, *p* = .010) (Figure 2B, Table S1B). In other words, Interrupted videos increased false memories in the Delayed group (β = -0.09, *t* = - 3.50, *p* < .001), but not the Immediate group (β = -0.01, *t* = -0.78, *p* = .437). We also found a main effect of group indicating more false memories in the Delayed group (β = -0.36, 95% CI [- 0.43, -0.29], *t* = -9.68, *p* < .001). Lastly, there was a main effect of reactivation type (β = -0.05, 95% CI [-0.08, -0.02], *t* = -3.31, *p* = .001), driven by the effect of prediction error increasing false memories in the Delayed group. To ensure that the effect of prediction error on memory updating was not driven by the first few trials (which are presumably most surprising), we also conducted a control analysis that found no effect of trial number on false memories (Supplementary Information, *Trial Number Control*, Table S2).

In sum, our behavioral results showed a novel dissociation between reinforcing and updating memories: Prediction errors during memory reactivation strengthened memories, evident both immediately and after a delay (Figure 2A). However, memory updating (as revealed by false memories) was not evident until after a delay (Figure 2B), consistent with prior studies that have been interpreted in terms of reconsolidation theory (1, 46–48).

#### Surprise Ratings and Semantic Similarity Predicted False Memories

Expanding on the results reported above, we recruited an independent sample to watch the videos and rate (on a 5-point Likert scale) the degree of surprise elicited by the narrative interruptions (Supplementary Information, *Supplementary Methods, Online Ratings of Stimulus Videos*). We then calculated the average surprise rating for each video and then related these surprise ratings to memory outcomes in our laboratory sample. Using linear mixed effects regression, we predicted subsequent *false memories* from the variables *reactivation type* (Full vs. Interrupted), *group* (Delayed vs. Immediate), *surprise ratings* (continuous), and all relevant interactions. All model parameters are reported in Table S3A. There was a significant interaction between surprise ratings and group (β = -0.03, 95% CI [-0.06, -0.01], *t* = -2.07, *p* = .039), such that more surprising videos were associated with more false memories selectively in the Delayed group (Delayed: β = 0.10, *z* = 2.49, *p* = .013; Immediate: β = 0.03, *z* = 0.95, *p* = .344). However, the three-way interaction among surprise ratings, group, and reactivation type was not significant (β = 0.01, 95% CI [-0.02, 0.04], *t* = 0.75, *p* = .451). Because our surprise ratings were collected from a separate online sample, this measure may not be sensitive enough to detect the expected three-way interaction when applied to the laboratory sample. In a separate model, we found that surprise ratings were not associated with correct details (Table S3B).

In the current study, we indexed memory updating in terms of false memories; however, incorporating relevant information into memory can be an adaptive function. We hypothesized that our paradigm would induce *false* memories because information would be integrated across semantically-related episodes. To test this hypothesis, we quantified semantic similarity among the 70 videos with a text-based analysis (Supplementary Information, *Supplementary Methods*, *Scoring of Memory Tests*). Using linear mixed effects regression, we predicted subsequent *false memories* from the variables *reactivation type* (Full vs. Interrupted), *group* (Delayed vs. Immediate), *semantic similarity* (continuous), and all relevant interactions. All model parameters are reported in Table S4. We found that videos that were more semantically-similar to other videos in the stimulus set produced more false memories (β = 0.11, 95% CI [0.04, 0.19], *t* = 3.09, *p* = .003) (Table S4). An interaction between semantic similarity and group predicted false memories (β = -0.04, 95% CI [-0.07, -0.01], *t* = -2.77, *p* = .006), such that the effect of semantic similarity was stronger in the Delayed group (β = 0.16, *z* = 3.90, *p* < .001) than in the Immediate group (β = 0.07, *z* = 1.80, *p* = .073). There was also an interaction between semantic similarity and reactivation type that predicted false memories (β = -0.03, 95% CI [-0.06, -0.01], *t* = -2.08, *p* = .038), such that the effect of semantic similarity was stronger for Interrupted videos (β = 0.15, *z* = 3.44, *p* < .001) than for Full videos (β = 0.08, *z* = 2.18, *p* = .030). However, the three-way interaction among reactivation type, group, and semantic similarity was not significant (β = 0.02, 95% CI [-0.02, 0.05], *t* = 0.99, *p* = .323). Overall, these results suggest that interfering details from semantically-related videos distorted memories; this interference effect was strongest for Interrupted videos and for participants in the Delayed group. Consistent with an adaptive updating process, memories may have been updated with relevant information from other videos.

### Univariate fMRI Results

The primary aim of our univariate fMRI analyses was to test the following questions: (1) Is hippocampal activation related to *reactivation type* (Full vs. Interrupted) and memory updating as indexed by subsequent *false memories?* (2) If so, does activation in the *basal forebrain* or the *VTA* moderate the relationship between hippocampal activation and memory updating?

We analyzed the blood oxygen level-dependent (BOLD) signal from the 24 subjects in the Delayed group (the Immediate group was not scanned). Our analyses focused on the fixation screen presented during the Post-Video period immediately after each video offset. This fixation screen was preceded by the narrative ending of each video, which was either as-expected (Full) or a surprising prediction error (Interrupted). Importantly, visual and auditory input was identical across conditions during this Post-Video fixation screen, and thus, by analyzing neural activation during the Post-Video period we controlled for differences in visual and auditory input (Figure 2B). Another possibility is that condition differences in the duration of visual stimulation (during video playback) could affect the magnitude and duration of the ongoing BOLD response during the Post-Video period. We conducted control analyses to confirm that our results were not confounded by video duration (Supplementary Information, *Video Duration Control*, Table S18).

#### The Effect of Hippocampal Activation on Memory Depended on Prediction Error

Whole-brain mass univariate results are provided in the Supplementary Information (*Whole-Brain Analysis*, Table S5, Figure S2). To investigate our primary research questions, we used single-trial modelling to relate post-video hippocampal activation to subsequent false memories. For our univariate analyses, we modelled a 2s impulse during the Post-Video period (fixation screen), convolved with the canonical double-gamma hemodynamic response function and phase-shifted 2s after video offset. This 2s shift targeted the peak Post-Video hippocampal response identified in previous studies (49, 50). We isolated BOLD activation during the Post-Video period on each trial and averaged parameter estimates across all voxels within each hippocampal ROI (Methods, *Univariate fMRI Analyses*).

Some past studies have shown that prediction error signals are stronger in left hippocampus and anterior hippocampus (18, 20, 21, 51), whereas posterior hippocampus is more sensitive to video offsets (52). Other studies have shown that anterior and posterior hippocampus parse continuous experience at different timescales (53, 54). On the basis of these findings, we tested separate (non-overlapping) ROIs for left anterior, right anterior, left posterior, and right posterior hippocampus (Supplementary Information, *Supplementary Methods, ROI Masks*). Activation estimates from these four ROIs are included within each model to test for left/right and anterior/posterior differences.

Using linear mixed-effects regression, we predicted trial-wise *hippocampal activation* from the following variables: *reactivation type* (Full vs. Interrupted), *false memories* (continuous measure), *hemisphere* (left vs. right), *axis* (anterior vs. posterior), and all relevant interactions. We found a significant interaction between reactivation type and subsequent false memories predicting hippocampal activation (β = -0.06, 95% CI [-0.09, -0.03], *t* = -4.33, *p* < .001) (Figure 3A) (Table S6). As predicted, this interaction demonstrated that the relationship between hippocampal activation and subsequent memory differed between conditions. After Full videos, greater hippocampal activation was associated with fewer subsequent false memories (Figure 3A, blue; β = -0.07, *z* = -2.81, *p* = .005), consistent with the idea that the hippocampus reinforces memory for episodes that just concluded (49, 50, 55). However, we observed the opposite effect when events were surprising. After Interrupted videos, greater hippocampal activation was associated *more* subsequent false memories (Figure 3A, orange; β = 0.05, *z* = 2.23, *p* = .026), consistent with the idea that surprise drives memory updating. Overall, this interaction demonstrated that the *same amount* of hippocampal activation predicted different memory outcomes depending on whether the video was Full (fewer false memories) or Interrupted (more false memories).

**Figure 3.**
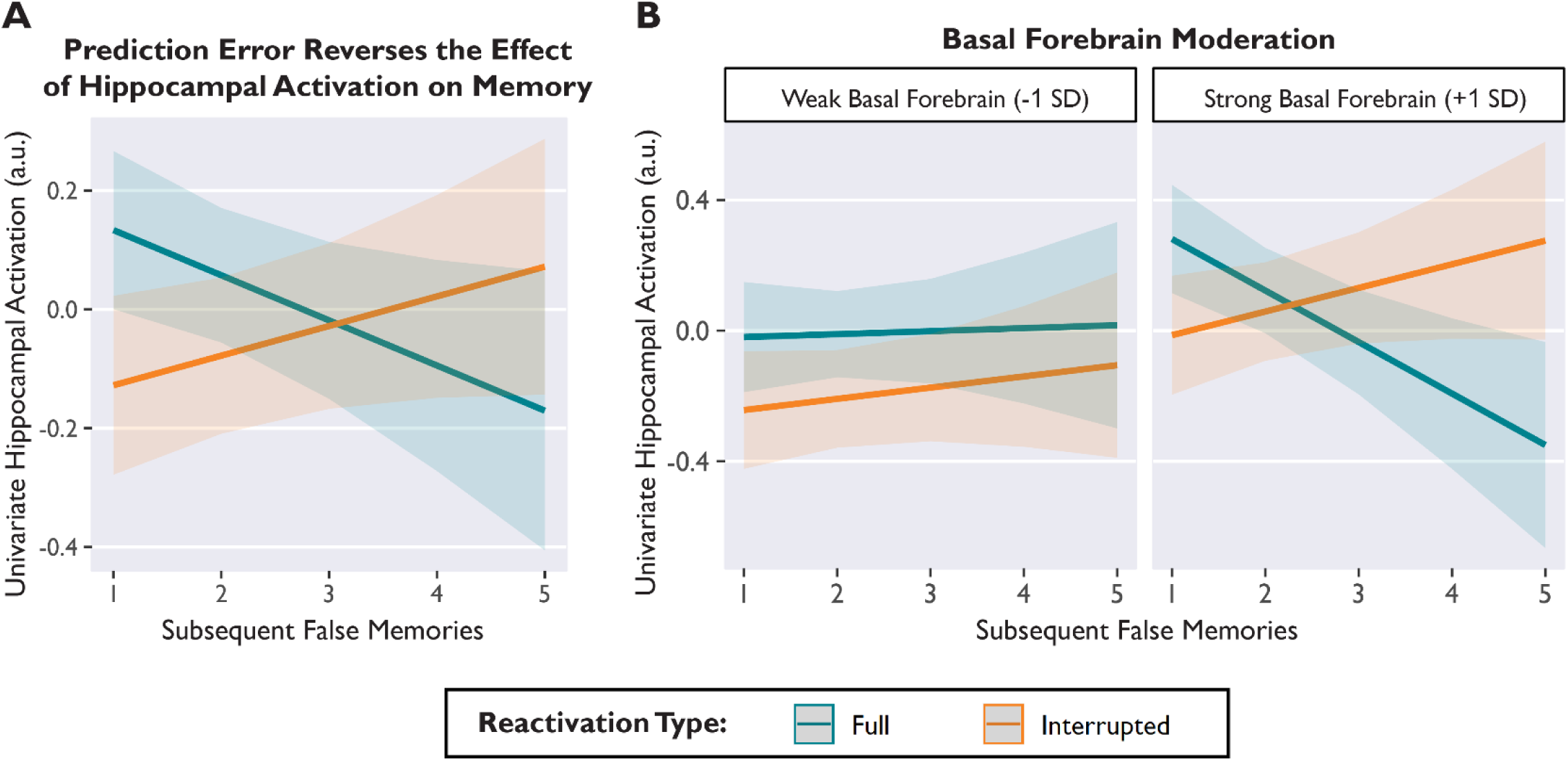
Prediction error reversed the relationship between average hippocampal activation (arbitrary units, a.u.) and subsequent memory, and this effect depended on concurrent basal forebrain activation. A) After Full-length videos, hippocampal activation was associated with memory *preservation*, predicting fewer false memories (blue). After Interrupted videos, hippocampal activation was associated with memory *updating*, predicting more false memories (orange). B) The effect of prediction error on hippocampal activation and memory was observed only when basal forebrain activation was strong (right). When basal forebrain activation was weak, hippocampal activation was unrelated to memory (left). Basal forebrain activation is binned (weak vs. strong) for visualization, but statistical models used a continuous variable. Lines depict model-predicted estimates, and shaded bands depict the 95% confidence interval. Model-derived estimates are shown instead of individual data points in order to show within-subjects effects, while controlling for subject and stimulus variance.

Neither the main effect of reactivation type on hippocampal activation (β = 0.04, 95% CI [-0.01, 0.09], *t* = 1.85, *p* = .069), nor the main effect of false memories on hippocampal activation (β = -0.01, 95% CI [-0.05, 0.03], *t* = -0.47, *p* = .640) were statistically significant. These null results demonstrate the value of examining the effect of prediction error on both hippocampal activation and memory outcomes. There was a main effect of axis indicating that average activation was greater in anterior hippocampus than posterior hippocampus (β = 0.04, 95% CI [0.01, 0.06, *t* = 3.12, *p* = .002). There was no main effect of hemisphere, and the axis and hemisphere variables did not interact with reactivation type or false memories. Parameter estimates for all variables are provided in Table S6.

Lastly, although our primary research questions pertained to false memories, we also tested a model that included both false memories and correct details (Table S7). We found that correct details also interacted with reactivation type to predict hippocampal activation (β = 0.03, 95% CI [0.01, 0.06], *t* = 2.10, *p* = .036), but numerically contrasted with our results for false memories. After Full videos, hippocampal activation was positively associated with correct details (β = 0.01, *z* = 0.33, *p* = .742), but was negatively associated with false memories (β = - 0.06, *z* = -2.44, *p* = .015). After Interrupted videos, these associations reversed direction: hippocampal activation was negatively associated with correct details (β = -0.05, *z* = -1.55, *p* = .120), but was positively associated with false memories (β = 0.05, *z* = 2.07, *p* = .039). Notably, although the overall interaction term was significant, the associations between hippocampal activation and correct details did not significantly differ from zero. Importantly, the effect of hippocampal activation on false memories remained unchanged after controlling for correct details (Full: β = -0.06, *z* = -2.44, *p* = .015; Interrupted: β = 0.05, *z* = 2.07, *p* = .039), demonstrating that the effect of memory updating was distinct from recall success for correct details (Table S7).

#### Hippocampal-Basal Forebrain Connectivity Predicted Memory Outcomes

Next, we tested hypotheses about neuromodulatory mechanisms by examining activation in the basal forebrain (which contains the medial septal nucleus, the primary source of acetylcholine in the hippocampus) (29, 31, 37) and the VTA (which contains dopaminergic neurons that project to the hippocampus) (16, 24, 56). First, we extracted average basal forebrain and VTA activation during the Post-Video period (i.e., the 2-second segment of the Post-Video fixation period, consistent with modelling of hippocampal activation).

Using linear mixed effects regression, we predicted *hippocampal activation* from the variables *reactivation type* (Full vs. Interrupted), *false memories* (continuous), *basal forebrain activation* (continuous), *hemisphere* (left vs. right), *axis* (anterior vs. posterior), and all relevant interactions. There was a significant three-way interaction among basal forebrain activation, reactivation type, and false memories predicting hippocampal activation (β = -0.04, 95% CI [- 0.06, -0.01], *t* = -2.66, *p* = .008) (Figure 3B). This interaction demonstrated that the relationship between hippocampal activation and subsequent memory (Figure 3A) was evident only when the basal forebrain was also strongly activated (Figure 3B, right) (Full: β = -0.12, *z* = -4.01, *p* < .001; Interrupted: β = 0.07, *z* = 2.37, *p* = .018). When basal forebrain activation was weak, hippocampal activation was unrelated to memory (Figure 3B, left) (Full: β = 0.01, *z* = -0.33, *p* = .739; Interrupted: β = 0.04, *z* = 1.39, *p* = .166). There was also a main effect of basal forebrain activation predicting hippocampal activation during the Post-Video period (β = 0.10, 95% CI [0.02, 0.19], *t* = 2.34, *p* = .029). Other results from this extended model were consistent with the base model (without basal forebrain variables) described above. There were no significant interactions with hemisphere or axis. All parameter estimates are provided in Table S8.

Next, we examined the role of VTA activation. We modified the model described above by replacing the basal forebrain activation variable (and interactions) with VTA activation parameters. VTA activation was positively related to hippocampal activation during the Post-Video period (β = 0.15, 95% CI [0.05, 0.25], *t* = 3.03, *p* = .006). However, there was no interaction among VTA activation, reactivation type, and false memories (β = -0.004, 95% CI [- 0.03, 0.02], *t* = -0.32, *p* = .747). Thus, there was no evidence that VTA moderated the effect of hippocampal activation on memory. All parameter estimates are provided in Table S9. In separate model, we also tested for a main effect of prediction error on basal forebrain activation (Table S10A) or VTA activation (Table S10B), and found no significant effects.

### Multivariate fMRI Results

Overall, our univariate results suggest that prediction error changed the role of the hippocampus: the same magnitude of hippocampal activation predicted opposing effects on memory depending on whether events were expected or surprising. Moreover, this effect of hippocampal activation on memory depended on concurrent basal forebrain activation, consistent with the idea that acetylcholine regulates hippocampal processing (27, 28). On the basis of our univariate findings, we proposed that during video playback, the hippocampus continually generates predictions and sustains episodic representations (57–59). If no prediction error is detected, these representations should be sustained, and the hippocampus should preserve and reinforce the memory (i.e., decreasing false memories). If a prediction error is detected, then the hippocampus should abandon ongoing predictions and prepare to update a memory (i.e., increasing false memories) (28, 29). Therefore, we hypothesized that prediction errors would disrupt sustained representations in the hippocampus, and that disrupting hippocampal representations would lead to memory updating. Furthermore, we predicted that activation in the basal forebrain and/or VTA would link hippocampal representations to memory outcomes, via neuromodulation of hippocampal processing (24–29, 34, 60, 61).

#### Prediction Errors Disrupted Sustained Representations in the Hippocampus

Past studies in rodents and humans have used *autocorrelation* measures, which quantify similarity across neural patterns, to investigate hippocampal representations during naturalistic tasks (53, 54). *Temporal autocorrelation* is an index of multivariate information that is preserved over time; this measures moment-to-moment overlap of activation patterns (53, 58, 62). Intracranial recordings in humans have shown that temporal autocorrelation in the hippocampus ramps up over the course of familiar episodes (58). Ramping autocorrelation reflects sustained neural representations, consistent with the hippocampus generating predictions and anticipating upcoming stimuli (57, 58). To test whether hippocampal representations were sustained or disrupted over time, we calculated temporal autocorrelation by correlating the activation of all voxels within the hippocampus at timepoint *T* with the activation pattern at timepoint *T+1 sec* (Methods, *Multivariate fMRI Analyses*). Importantly, we also investigated autocorrelation in two control regions (inferior lateral occipital cortex and white matter) to demonstrate that these representational changes were not a spurious whole-brain phenomenon (Supplementary Information, *Autocorrelation Control*).

First, we tested whether autocorrelation increased during video playback (binned into 5-second *video segments* over time). We used linear mixed effects regression to predict *hippocampal autocorrelation* (averaged over 5s bins) from the variables *video segment* (continuous)*, hemisphere, axis,* and all interaction terms. We found that hippocampal autocorrelation increased linearly as videos progressed (β = 0.025, 95% CI [0.01, 0.04], *t* = 3.41, *p* = .002), suggesting that episodic representations were sustained and stabilized during video playback (Figure 4A) (58). There were no significant effects of hemisphere or axis. All parameter estimates are provided in Table S11A.

**Figure 4.**
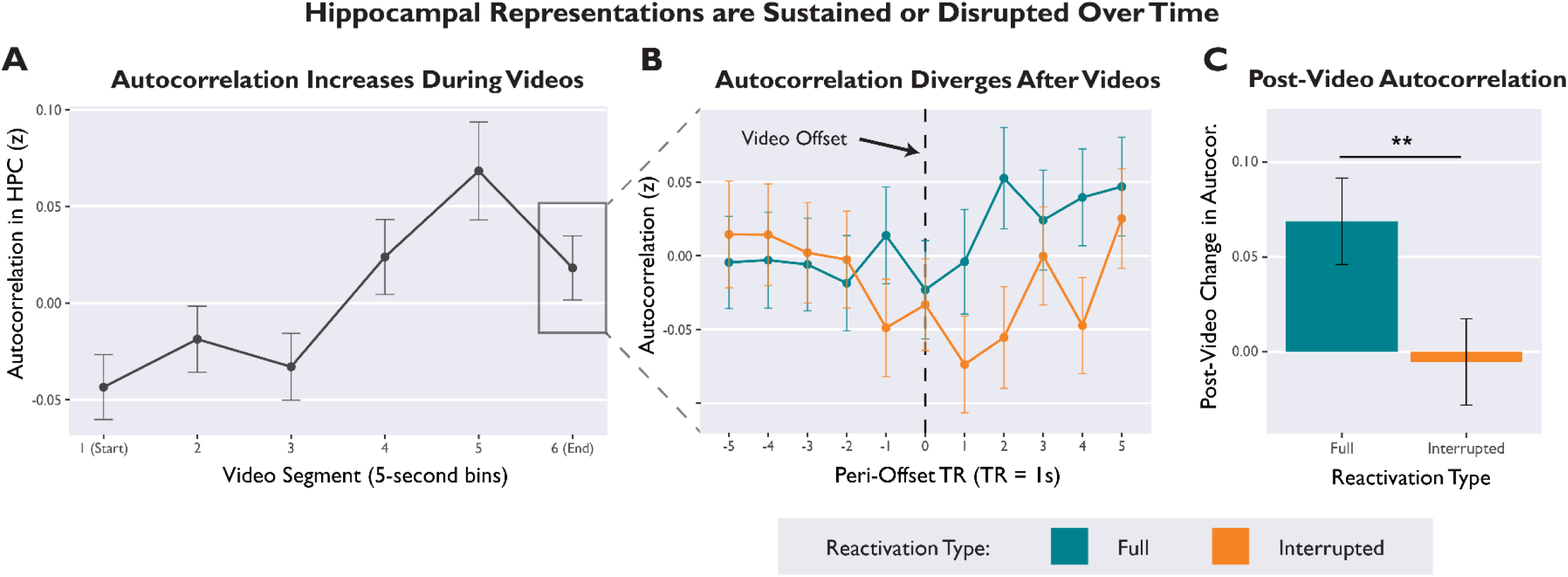
Hippocampal representations are sustained or disrupted over time, depending on whether or not episodes align with expectations. A) Temporal autocorrelation in the hippocampus gradually increased over the course of a video, suggesting that episodic representations were sustained over time. Autocorrelation values were averaged over 5-second bins of video playback. B) Autocorrelation trajectories for Full and Interrupted videos diverged during the Post-Video period. Plot visualizes second-by-second autocorrelation values in the hippocampus, time-locked to the moment of video offset (black dotted line). C) Average Post-Video change in autocorrelation (average autocorrelation scores for the 5-sec bin immediately *after* video offset, minus average autocorrelation for the bin immediately *before* offset). Hippocampal representations were sustained after Full videos, but disrupted after Interrupted videos. Error bars depict SEM.

Next, we tested whether prediction error disrupted this ramping autocorrelation. To analyze Post-Video *change in autocorrelation*, we calculated a difference score for each trial by subtracting the average autocorrelation value from the 5s bin immediately *before* video offset from the average autocorrelation value from the 5s video immediately *after* video offset. We then used linear mixed effects regression to predict average *change in autocorrelation* from the variables *reactivation type*, *hemisphere*, *axis*, and interaction terms. There was a significant main effect of reactivation type (β = 0.04, 95% CI [0.01, 0.07], *t* = 2.21, *p* = .038), such that autocorrelation increased after the offset of Full videos but not Interrupted videos (Figure 4C). In other words, prediction errors disrupted the continuity of hippocampal representations. This Post-Video divergence is visualized in Figure 4B. There were no significant interactions with hemisphere or axis. All parameter estimates are provided in Table S11B.

#### Disruption of Hippocampal Autocorrelation Predicted False Memories

Next, we tested whether disruption of hippocampal representations predicted memory updating. Using linear mixed effects regression, we predicted subsequent *false memories* from the variables *reactivation type*, Post-Video *change in autocorrelation* (continuous), *hemisphere*, *axis*, and all relevant interactions. We also included a continuous covariate for univariate hippocampal activation (thus controlling for any autocorrelation effects that may be a consequence of univariate activation). There was a significant interaction between reactivation type and change in autocorrelation predicting false memories (β = 0.04, 95% CI [0.02, 0.07], *t* = 3.34, *p* < .001) (Figure 5A). After Interrupted videos, disrupting hippocampal representations led to memory updating (β = -0.06, *z* = -2.89, *p* = .004). Conversely, after Full videos, hippocampal autocorrelation was not related to false memories (β = 0.02, *z* = 1.03, *p* = .303). There were no significant interactions with hemisphere or axis. All other parameters are reported in Table S12A.

**Figure 5.**
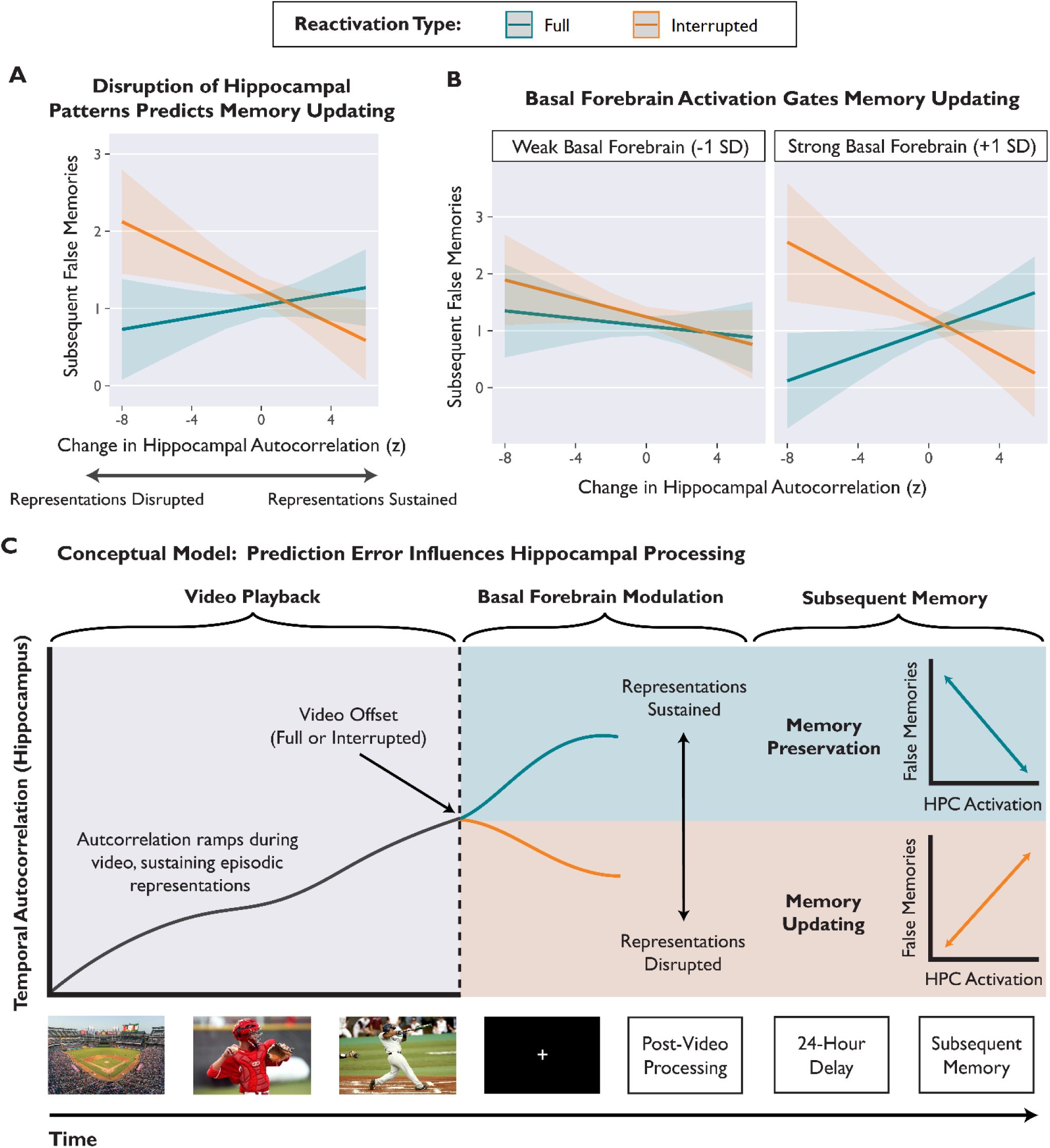
Prediction errors elicited by Interrupted videos disrupted sustained hippocampal representations, and this disruption predicted memory updating. A) Estimated values from a linear regression model predicting subsequent false memories from the interaction of reactivation type and change in autocorrelation. After Interrupted videos, decreases in autocorrelation were related to increased memory updating. B) The effect of prediction error on hippocampal autocorrelation and subsequent memory depended on concurrent basal forebrain activation. Basal forebrain activation was binned (weak vs. strong) for visualization, but statistical models used a continuous variable. Shaded bands depict 95% confidence intervals around the regression line. In panels A and B, model-predicted estimates are depicted instead of individual data points in order to show within-subject effects, while controlling for subject and stimulus variability. C) Conceptual schematic depicting the effect of prediction error on hippocampal representations and subsequent memory. During a video, the hippocampus sustains episodic representations over time, consistent with generating ongoing predictions. After video offset, the hippocampus acts to preserve the memory (representations sustained) or prepare for memory updating (representations disrupted). The link between prediction error and memory outcomes depends on co-activation of the hippocampus and basal forebrain during the Post-Video period.

#### Basal Forebrain Activation Links Hippocampal Autocorrelation to Memory

What determines whether hippocampal representations are sustained or disrupted? To investigate candidate neuromodulatory mechanisms, we extended the model described above by adding parameters for average Post-Video basal forebrain activation and interaction terms. The model included all relevant interaction terms, reported in full in Table S12B. Paralleling our univariate findings, we found a significant three-way interaction among basal forebrain activation, reactivation type, and hippocampal autocorrelation that predicted subsequent false memories (β = 0.03, 95% CI [0.01, 0.06], *t* = 2.54, *p* = .011) (Figure 5B). In other words, prediction errors disrupted hippocampal representations and led to memory updating, but only when the basal forebrain was also strongly activated (at +1SD basal forebrain activation, Interrupted: β = -0.07, *z* = -2.46, *p* = .014; Full: β = 0.08, *z* = 2.92, *p* = .004) (Figure 5B, right).

Next, we tested whether VTA activation was related to hippocampal autocorrelation and memory. We modified the model described above (predicting subsequent false memories) by replacing the basal forebrain activation variable (and interaction terms) with the VTA activation variable. In contrast to our basal forebrain findings, there was no three-way interaction among VTA activation, reactivation type, and hippocampal autocorrelation (β = -0.01, *t* = -0.54, *p* = .587, 95% CI [-0.03, 0.02]). All parameter estimates are reported in Table S12C. Overall, we found that our autocorrelation results paralleled our univariate findings: basal forebrain activation, but not VTA activation, was crucial for connecting hippocampal representations to memory outcomes.

## Discussion

Here, we show that prediction errors modulate the function of the hippocampus and allow memories to be modified, consistent with an adaptive updating mechanism. In our fMRI paradigm, we elicited mnemonic prediction errors by interrupting familiar narrative videos immediately before the expected conclusions. Prediction errors reversed the relationship between univariate hippocampal activation and subsequent memory: After expected video endings, hippocampal activation was associated with memory preservation, but after prediction errors, hippocampal activation was associated with memory updating. Tracking the stability of hippocampal representations revealed that prediction errors disrupted activation patterns; this pattern disruption predicted memory updating. Crucially, the association between hippocampal activation (both univariate and multivariate) and memory outcomes depended on concurrent basal forebrain activation during the Post-Video period. We conclude that prediction error, coupled with basal forebrain modulation, prompts the hippocampus to abandon ongoing predictions and prepare to update a memory by incorporating other information present in the environment (Figure 5).

### Prediction Errors Disrupt Hippocampal Representations and Update Memories

Past studies of mnemonic prediction errors have reported an increase in univariate hippocampal activation, but have not examined whether this neural signal affects memory (17–19, 63). For the first time, we show that after prediction errors, hippocampal activation leads to memory updating. Crucially, we demonstrate that univariate measures are *insufficient* for understanding the effect of prediction error on the hippocampus, because the same amount of hippocampal activation can exert opposing effects on memory. Prediction error reversed the relationship between hippocampal activation and subsequent memory, suggesting a shift in hippocampal processing (Figure 3). After expected endings (Full videos), hippocampal activation protected against false memories, consistent with the idea that the hippocampus reinforces memory after the conclusion of an episode (49, 50). In contrast, after surprising endings (Interrupted videos), hippocampal activation predicted *more* false memories, consistent with the idea that prediction errors can destabilize memories and enable updating (1–3). Overall, this reversal supports the idea that the hippocampus acts to *preserve* memories after expected events, but *update* memories after surprising events. These divergent modes of processing parallel prior studies on encoding vs. retrieval modes (20, 27, 64) and internal vs. external attention (28).

To test the idea that prediction errors influence hippocampal processing, we tracked hippocampal activation patterns to examine how episodic representations were sustained or disrupted over time. We used *temporal autocorrelation* (the moment-to-moment overlap of activation patterns) as a measure of continuity in hippocampal representations (53, 58, 62). As narratives progressed, autocorrelation increased, reflecting stability and continuity; this increase in autocorrelation suggested that the hippocampus generated predictions (57, 58) and sustained episodic representations over time (49, 65) (Figure 4A). Crucially, prediction errors disrupted the stability of hippocampal representations (Figure 4B, 4C), and this disruption predicted the degree of memory updating (Figure 5A). Overall, we propose that disruption of hippocampal representations indicates a shift in processing: Prediction error prompts the hippocampus to abandon ongoing predictions and prepare to update a memory (Figure 5C).

Our univariate findings also diverge from past studies of hippocampal prediction error responses, which have shown that hippocampal activation increases after prediction errors (17–19, 63). In contrast, our single-trial analyses showed no significant main effect of reactivation type on hippocampal activation; the effect of prediction error was revealed only when examining the link between hippocampal activation and subsequent memory. Moreover, our whole-brain mass univariate analyses revealed a significant cluster in left hippocampus that was *less* activated after Interrupted videos than Full videos (Supplementary Information, *Whole-Brain Analysis*). Our task elicits surprise by omitting expected endings, comparable to a *negative* prediction error. Previous studies have elicited surprise by replacing expected stimuli with novel stimuli, thus adding new information, comparable to a *positive* prediction error. Therefore, this reversal of activation (Full > Interrupted trials) is consistent with prior studies of reward (13) and information prediction errors (4, 66), which have shown *increases* in neural activation after positive prediction errors and *decreases* after negative prediction errors. Overall, our results suggest that hippocampal responses may depend on the stimuli and type of surprise, and demonstrate that prediction errors change the effect of hippocampal activation on memory.

### Basal Forebrain Activation Relates to Hippocampal Processing

Past studies have suggested that either cholinergic (27–29, 35–37, 60) or dopaminergic (16, 24, 61) modulation could regulate hippocampal function after prediction errors, such as by enhancing plasticity and switching between processing modes. However, mixed evidence supporting both hypotheses has left the question unresolved (25, 26, 31, 33, 34). Here, we investigated whether activation of the basal forebrain or the VTA could explain the relationship between hippocampal activation after prediction error and subsequent memory. We found that the effect of prediction error on memory depended on co-activation of the hippocampus and basal forebrain, suggesting that connectivity between these regions is important for shifting hippocampal processing modes to either preserve or update a memory. Hippocampal activation was associated with memory updating after prediction errors, but only when the basal forebrain was also activated. Likewise, disrupting hippocampal representations led to memory updating after prediction errors, but only when the basal forebrain was also activated. Although fMRI cannot provide direct evidence of neuromodulation, our results are consistent with the idea that cholinergic modulation from the basal forebrain (27, 35–37, 39) influences hippocampal processing and memory outcomes.

Our findings are also relevant to the functional relationship between the VTA and hippocampus. Although we found a robust positive correlation between VTA and hippocampal activation during the Post-Video period, VTA activation was unrelated to prediction error and did not link hippocampal activation with memory outcomes. These findings are consistent with our prior proposal that connectivity between the VTA and hippocampus reflects modulation of hippocampal learning states by sustained VTA activity (16, 69) rather than phasic VTA responses (25, 70–72). However, our paradigm was optimized for detecting memory updating instead of midbrain prediction error responses. It is also possible that prediction error signals could be transmitted to the hippocampus from the locus coeruleus (71, 72). Analyses of locus coeruleus in the current data did not reveal any relationships. Future research could disambiguate the roles of the basal forebrain, VTA, and locus coeruleus by examining both event-related and sustained connectivity with the hippocampus and their consequences for memory.

### Prediction Error Both Strengthens and Updates Memories

Comparing behavioral results across the Delayed and Immediate groups revealed a dissociation: prediction error both strengthened and updated memories, but over distinct timecourses and likely via different mechanisms (Figure 2). Prediction error increased the number of correct details recalled, both immediately and after a one-day delay. This finding is consistent with recent evidence that mnemonic prediction errors can promote detailed memories immediately (73), possibly by way of enhanced attention and pattern separation. In contrast, we found that prediction error increased false memories only after a delay, consistent with prior studies that have shown that memory updating requires a delay for protein synthesis to occur (1, 47, 74). Although it is possible that the relatively few false memories in the Immediate group made it more difficult to detect an effect of prediction error, a control analysis showed that excluding low-variance subjects from the Immediate group did not change our results (Supplementary Information, *Behavioral Variance Control*). Overall, our finding that prediction error increased both correct details and false memories supports the idea that surprise drives adaptive updating.

In the present study, we used false memories as an index of memory updating. In the real world, however, memory updating can be adaptive: new information is not “false” *per se*, but a relevant addendum to prior knowledge. In our paradigm, interference from other stimulus videos likely produced false memories because information was integrated across videos. Previously, we found that prediction errors selectively updated memories with semantically-related information from new videos that were specifically chosen to interfere with reactivated memories (48). Here, we showed that videos that shared greater semantic similarity with the rest of the stimulus set produced more false memories (Table S4). This finding suggests that prediction error increases source confusion or integration of information across related memories. Prior studies have shown that memory updating occurs when interference is introduced after a memory trace is reactivated (1, 47, 75). In our paradigm, interference could arise from semantically-related details that result from the subject recalling related memories, or visual input from subsequent videos during the task. This finding accords with prior behavioral studies (1, 47, 75) and computational models of event segmentation (76, 77), which have both shown that interference among related episodes can produce false memories and/or source confusion after prediction error. However, memory updating is beneficial in other situations that require integrating old and new knowledge, or correcting erroneous information.

One possible mechanism for the memory updating we observed is *reconsolidation*, the process by which reactivating a memory trace can temporarily render it malleable. However, evidence for cellular reconsolidation processes in humans is lacking. Although previous human studies have used reconsolidation-like paradigms to demonstrate memory malleability (4, 47, 48, 75), it remains unknown whether the synaptic mechanisms of reconsolidation are consistent across animals and humans (1). A key prediction of reconsolidation theory is that memory updating effects should only emerge after a delay, because the process of modifying and restabilizing a memory trace requires protein synthesis that occurs over several hours. We found that the behavioral effect of prediction error on memory updating required a delay, which aligns with this theoretical prediction. Overall, our findings are broadly relevant to research on prediction error and memory, and reconsolidation theory offers one possible framework.

### Limitations and Future Directions

We were unable to directly test whether the relationship between neural activation and memory updating required a delay, because the fMRI participants always completed the delayed memory test and may show effects of memory updating as well as forgetting over time. Although our prior behavioral work demonstrated an effect of prediction error that was not confounded with the delay (48), scanning a group of participants who experience a delay-to-test without memory reactivation would enable an investigation of the neural correlates of immediate vs. delayed memory effects.

In the present study, we elicited surprise by interrupting videos before their expected narrative endings. However, we were not able to directly measure subjective surprise, because asking participants to rate surprise after each video would have disrupted other cognitive processes and revealed the goal of the manipulation. To ensure that Interrupted videos would continue to elicit surprise after the first few trials, we included the following features in our experimental design: 1) Full-length videos were presented twice during encoding to set strong expectations; 2) Interrupted videos violated strongly-expected action-outcome contingencies (e.g., a baseball batter halted mid-swing); 3) trials were pseudorandomized so that participants could not anticipate whether each video would be Full or Interrupted; and 4) participants did not know *when* each video would be interrupted. Consistent with these design considerations, we found that the effect of prediction error on false memories did not interact with trial number, suggesting that surprise did not diminish over the course of the experiment (Table S2).

Prior studies with humans and animals have used *incomplete reminders* (e.g., a conditioned stimulus without the expected outcome) to elicit prediction error (1, 3, 75). Here, we mimicked this approach by interrupting narrative videos and omitting the expected endings. Incomplete reminders may be particularly effective because they elicit memory reactivation, and memory reactivation supports plasticity (78–80). After Interrupted videos, participants may actively retrieve the missing endings; this memory reactivation could contribute to memory malleability. As discussed in our prior review (1), both prediction error and memory reactivation may interact to support memory updating. Memory reactivation may be a prerequisite for generating a prediction, and experiencing a prediction error may prompt further memory reactivation. Future studies could directly investigate the role of memory reactivation by asking participants to report their ability to recall the missing endings, or by testing encoding-retrieval pattern similarity.

Lastly, a limitation of the present study is that we were unable to determine the temporal distribution or the source of each individual false memory. Most details that participants recalled (both correct and incorrect) pertained to the entire video, such as perceptual details about the characters and setting. Many of these perceptual details were also shared across multiple videos (e.g., several characters with green shirts), making it impossible to determine the source of each specific detail. Interestingly, we found that the omitted endings on Interrupted trials were rarely associated with false memories. Recall of the narrative content of the video endings was very accurate: in the Delayed group, only 2% of all false memories specifically pertained to the missing ending. Because the salient action-outcome contingencies define the videos, even when interrupted, these central details were not likely to be affected. Instead, we found that prediction error was more likely to induce a holistic distortion of perceptual content from throughout each video (e.g., details about the setting or a character).

## Conclusion

The brain continually generates predictions based on past experiences. When expectations do not align with reality, memories should be updated with relevant new information. We propose that mnemonic prediction errors prompt the hippocampus to abandon ongoing predictions and update memories by incorporating relevant details from subsequent experiences. In this way, surprise modulates hippocampal processing and determines the fate of episodic memories. This theoretical framework of memory updating bears implications for eyewitness testimony, education, and treating pathological memories (e.g., in Post-Traumatic Stress Disorder). Beyond memory research, our results offer new insights for theories on the whole-brain predictive processes that govern attention, perception, action, and decision-making.

## Acknowledgements

This research was funded by grants awarded to MDB from the *James S. McDonnell Foundation* (Scholar Award in Understanding Human Cognition) and the *Natural Sciences and Engineering Research Council* of Canada (Discovery Grant and Accelerator Supplement, RGPIN-2014-05959 and RGPIN-2020-05747). AHS has been supported by awards from the *National Science Foundation* (Graduate Research Fellowship) and *Natural Sciences and Engineering Research Council* of Canada (Postgraduate Doctoral Scholarship, Undergraduate Student Research Award). We also give thanks to Carolyn Chung, Tolulemi Gbile, and Aria Fallah for their invaluable contributions to data collection, transcription, and scoring. We thank Jia-Hou Poh for helpful comments on the manuscript.

## Author Contributions

AHS and MDB developed the study design. AHS programmed the study, collected data, conducted analyses, and drafted the manuscript. GMM contributed substantially to data collection and IKB contributed to autocorrelation analyses. MDB and RAA contributed to the analysis approach and interpretation of results. All authors contributed to revising the manuscript and approved the final version.

## Methods

### Data, Code, and Materials

Brief descriptions of the stimulus videos are provided in Table S13. The full set of stimulus videos, along with derivative data and code necessary to reproduce results, are provided online in the project repository hosted by the Open Science Framework (https://osf.io/xb7sq/).

### Participants

We recruited 55 paid participants from the University of Toronto community (Delayed group: $70, Immediate group: $40). Seven participants were excluded (see Supplementary Information, *Exclusions*), yielding a final sample of 48 participants. The sample size was determined *a priori* to satisfy the following conditions: (1) reproduce the sample size and achieve at least 90% power to detect the interaction effect found in a prior study (*η_p_^2^* = 0.17) (48), and (2) evenly allocate participants to 6 pseudorandomized trial order lists. Participants were healthy young adults (age: *M =* 22.42, *SD =* 2.41, range [18, 30]; gender: 75% female, 25% male). Inclusion criteria were as follows: between the ages of 18-30, normal or corrected-to-normal vision and hearing, no history of neurological or psychiatric disorders, and fluency in English. fMRI participants were all right-handed. All participants provided informed consent, and the study was approved by the University of Toronto Institutional Review Board, Protocol #00035787.

In consideration of the effects of sleep on consolidation, we also asked participants to report approximate hours of sleep over the course of the study. Participants slept an average of 7.28 hours (*SD* = 1.31) between the Day 1 and Day 2 sessions, and Delayed group participants slept an average of 7.02 hours (*SD* = 1) between the Day 2 and Day 3 sessions.

### Stimuli

Stimulus videos were sourced from movies, TV, and YouTube clips. We chose 70 videos that featured distinct narrative events (duration *M* = 30 sec, *SD* = 7 sec). Semantic similarity varied across videos (e.g., several videos featured sporting events), but there were no overlapping scenes or characters. During pilot testing, we ensured that the videos would be infrequently recognized by our participants. The 70 videos used in the experiment are described in Table S13 and publicly available on the Open Science Framework (https://osf.io/xb7sq/). The Interrupted version of each video ended abruptly at the narrative climax, omitting the salient ending and violating expectations (duration *M =* 25 sec, *SD =* 4 sec).

### Procedure

During the **Encoding session**, participants viewed all 70 videos in full-length form (randomized order). Each video was presented twice in a row to ensure that participants had strong expectations about the narrative outcomes for each video, a prerequisite for eliciting prediction error later.

During the **Reactivation session**, participants viewed each video again a single time (35 Full videos, 35 Interrupted videos). Videos were played in a pseudorandom order (six trial order lists, counterbalanced across participants) such that there were never more than two consecutive Interrupted videos. Participants could not reliably anticipate whether each video would be Interrupted, or where the interruption might occur. We also performed eye-tracking during the Encoding and Reactivation sessions for participants in both the Delayed and Immediate groups (EyeLink v.1000+, SR-Research). Eye-tracking was used to monitor alertness during the task, but these data are not discussed further.

Lastly, the **Test session** involved a structured interview-style recall test about details from each of the videos. Participants were cued with the name of each video and prompted to recall the narrative. The experimenter then probed the participant for more information with a pre-determined list of questions (e.g., “Can you describe the setting or context of the video?”, “Can you describe what the character looked like? Do you remember gender, age range, hair color, or clothing?”). Participants were instructed to answer based on their memory of the Full-length videos that had been originally presented during encoding. Because we were interested in false memories as a measure of memory modification, we instructed participants not to guess and permitted them to skip details they could not recall.

Overall, the experiment took place over three days for participants in the Delayed group (24-hour delays between Encoding, Reactivation, and Test), or over two days for participants in the Immediate group (24-hour delay between Encoding and Reactivation, no delay between Reactivation and Test). Only the Delayed group underwent neuroimaging.

Consistent with past studies (81–83), we maintained consistent contextual factors between Encoding, Reactivation, and Test sessions. Delayed group participants completed the encoding session in a mock scanner (shell of a retired 1.5T Siemens Avanto scanner), while recorded MRI sounds were played in the background. The mock scanner room was adjacent to the real scanner room, and was very similar in size and appearance. Delayed group participants completed the Reactivation session in the real fMRI scanner and the Test session at a desk in the mock scanner room. Participants in the Immediate group completed all three sessions in the same behavioral testing room. In both groups, participants completed all three sessions with the same experimenter.

### Statistical Analysis

For both behavioral and neural data, we conducted trial-wise analyses with linear mixed-effects regression models. Detailed information about the random effects specification is provided in the Supplementary Information (*Supplementary Methods, Linear Mixed-Effects Regression*).

#### Univariate fMRI Analyses

Whole-brain mass univariate results are reported in the Supplementary Information (*Whole-Brain Analysis*, Figure S2, Table S5). Details about preprocessing steps and region-of-interest masks are provided in the Supplementary Information (*Supplementary Methods, fMRI Preprocessing, Region of Interest Masks*). The primary findings reported in the main text reflect a single-trial modelling approach that estimated hippocampal responses to each video during the task. In order to isolate responses on each trial, we employed the Least Squares-Single approach and constructed a separate GLM for each trial (84, 85). We modelled each trial as a 2s impulse in the Post-Video period, convolved with the canonical double-gamma hemodynamic response function and phase-shifted 2s after video offset. This 2s shift targets the peak hippocampal response previously identified in studies of post-video processing (49, 50). Within each GLM, the target trial (2s event) was isolated as one regressor, and all other events were modelled with a separate regressor for each type of event (e.g., video playback, video name cues, other fixation periods). For each trial, we masked the processed data and averaged across voxels within each ROI to yield an average activation value.

#### Multivariate fMRI Analyses

Multivariate temporal autocorrelation analyses (53, 58) were conducted on preprocessed data (prior to single-trial GLM analysis). We extracted the whole-run timeseries from every voxel within each ROI using the *fslmeants* utility. For control analyses (white matter and iLOC ROIs), autocorrelation was calculated on 200 contiguous voxels, approximately matching the size of the hippocampal ROIs. Temporal autocorrelation was defined as the Pearson product-moment correlation between all voxel activation values at timepoint T and timepoint T+1s. This method yielded an autocorrelation value for every second of each functional run, excluding the final TR. Autocorrelation values were standardized (Fisher’s *z*) prior to statistical analysis.

Next, we aligned multivariate timeseries data with event onset and duration markers. Comparable to past research, we phase-shifted the timeseries by 4 seconds in order to account for HRF lag (86). This manual shifting is necessary because event onset regressors have not yet been convolved with the HRF (unlike in standard GLM analyses used for our univariate analyses). Note that our univariate analyses involved an additional 2s shift in addition to the standard HRF shift; this allowed us to target the peak post-video hippocampal response on each trial, but was not necessary for the autocorrelation analyses that yield TR-by-TR values.

After alignment, we calculated average autocorrelation values that were time-locked to events. For statistical analyses, autocorrelation values were averaged across 5-second bins during and after each video. To analyze signal history over the course of video playback, we related the video segment number (5s bins) to average autocorrelation values. For each video, we included the first five seconds (timepoints 0-4), the next four middle segments (timepoints 5-9, 10-14, 15-19, and 20-24), and the last five seconds (variable depending on the length of the video). This binning scheme spans the average video length of 30 seconds; additional middle segments from videos that were longer than 30 seconds were omitted. Lastly, to analyze Post-Video changes in autocorrelation, we calculated trial-by-trial difference scores by subtracting the average autocorrelation value for the 5-second bin immediately *before* video offset from the average autocorrelation value for the 5-second bin immediately *after* video offset. Autocorrelation values and difference scores for each trial were then submitted to linear mixed effects regression.

