## Supplementary Information for "Prediction Errors Disrupt Hippocampal Representations and Update Episodic Memories"

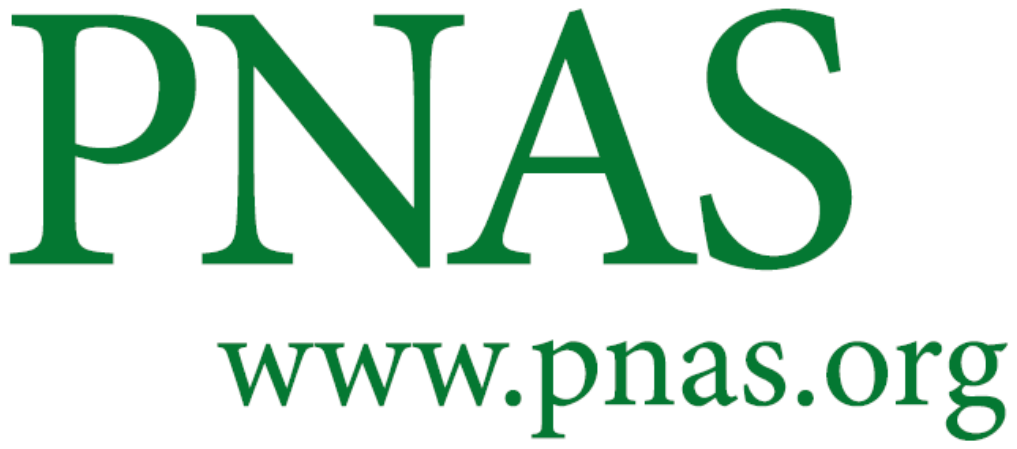

**Supplementary Information for**

Prediction Errors Disrupt Hippocampal Representations
and Update Episodic Memories

*Alyssa H. Sinclair, Grace M. Manalili, Iva K. Brunec, R. Alison Adcock, & Morgan D. Barense*

Corresponding Author: Alyssa H. Sinclair

**This PDF file includes:**

Supplementary text

Figures S1 to S3

Tables S1 to S18

SI References

**Other supplementary materials for this manuscript include the following:**

Data and code are provided in an OSF repository: https://osf.io/xb7sq/

**Please refer to the table of contents on the following page for further information.**

**Supplementary Information: Table of Contents**

| **Section/Item** | **Description** | **Page** |
| --- | --- | --- |
| Supplementary Methods | fMRI Scanning, Scoring of Memory Tests, Online Ratings of Stimulus Videos, Exclusions, Linear Mixed-Effects Regression, fMRI Preprocessing, Region of Interest Masks | 3-5 |
| Table S1 | Behavioral analyses predicting false memories and correct details | 6 |
| Trial Number Control,  Table S2 | Control analysis testing whether the effect of reactivation type on memory diminishes over the course of the experiment | 7 |
| Confidence & Forgetting, Figure S1 | Behavioral analyses for confidence ratings and forgotten videos, with plots of averages by condition | 8 |
| Table S3 | Behavioral analyses testing the effect of surprise ratings | 9 |
| Table S4 | Behavioral analyses testing the effect of semantic similarity | 10 |
| Behavioral Variance Control | Control behavioral analysis investigating variance by Group | 11 |
| Whole-Brain Analysis | Summary of whole-brain mass univariate results | 12 |
| Figure S2 | Visualization of whole-brain results, depicting Full > Interrupted and Interrupted > Full contrasts | 12 |
| Table S5 | Coordinates and statistics for significant clusters from the whole-brain mass univariate analysis | 13 |
| Table S6 | Neural analysis predicting univariate hippocampal activation from reactivation type and false memories | 14 |
| Table S7 | Neural analysis predicting univariate hippocampal activation from reactivation type, false memories, and correct details | 15 |
| Table S8 | Neural analysis predicting univariate hippocampal activation from reactivation type, false memories, and basal forebrain activation | 16 |
| Table S9 | Neural analysis predicting univariate hippocampal activation from reactivation type, false memories, and VTA activation | 17 |
| Table S10 | Neural analysis predicting basal forebrain (A) and VTA (B) activation from reactivation type and false memories | 18 |
| Table S11 | Neural analyses predicting hippocampal autocorrelation during and after videos | 19 |
| Table S12 | Neural analyses predicting subsequent false memories from hippocampal autocorrelation and univariate activation in BF/VTA | 20-23 |
| Table S13 | Brief descriptions of all stimulus videos | 23-24 |
| Tables S14-S16 | Parameter estimates from models predicting memory outcomes from video memorability, valence, and intensity | 25-27 |
| Basal Forebrain SNR, Table S17 | Alternative basal forebrain results, excluding the 5 subjects with lowest tSNR in the ROI. Compare to Table 11B. | 27-29 |
| Autocorrelation Control | Analysis of post-video autocorrelation in control regions | 30 |
| Video Duration Control, Table S18 | Control analyses that rule out video duration as a confounding variable | 30-32 |
| SI References | Sources for content cited in the Supplementary Information | 33 |

**Supplementary Methods**

**fMRI Scanning**

Scanning was performed with a 3T Siemens Prisma MRI scanner located at the Toronto Neuroimaging Center, University of Toronto. High-resolution functional images were collected with a T2*-weighted multiband-accelerated echo-planar imaging (EPI) pulse sequence, and a 32-channel head coil. Foam padding was used to minimize head motion. We acquired whole-brain BOLD activation estimates with a spatial resolution of 2.7mm isotropic voxels (TR: 1000 ms, TE: 29 ms, flip angle: 50**°**, 60 slices at transversal orientation, phase encoding: A>P, FoV: 210mm, Partial Fourier: 0.875, multiband factor: 4). High resolution T1-weighted anatomical images were acquired with a magnetization-prepared rapid-acquisition gradient-echo (MP-RAGE) pulse sequence (voxel size: 1mm isotropic, TE: 24 ms, TR: 2000 ms, TI: 1100 ms, flip angle: 9**°**) to allow 3D reconstruction and volume-based statistical analysis.

For the fMRI version of the task (Delayed group), stimuli were presented with EyeLink Experiment Builder (SR-Research) on a BOLDscreen display monitor (32”, 1920x1090, 100Hz refresh rate), viewed through a mirror attached to the head coil. Auditory stimulation was presented with in-ear MRI-compatible headphones (Sensimetrics, model S14). During the initial scout scan, we performed a sound test by playing the soundtrack of a movie trailer (not included in the stimulus videos) and adjusting the volume. For the behavioral version of the task (Immediate control group), videos were presented on a desktop computer and audio was presented with over-ear headphones.

**Scoring of Memory Tests**

We transcribed memory tests with *Temi*, an automated voice-to-text tool, then manually edited transcripts to verify accuracy (https://www.temi.com/). We coded videos as “forgotten” if the participant entirely failed to retrieve a memory when cued with the name of the video and a hint from a pre-determined list (brief descriptions of each video, provided in Table S13). Scoring of details was conducted with *NVivo 12*, a program for qualitative analysis of transcripts. Research assistants manually labelled each detail as correct or false. Scorers were blinded to subject identity and reactivation type (Full vs. Interrupted) while scoring the memory tests. The number of false memories per-trial ranged from 0-6, but there were very few trials with 5 or 6 false memories. To account for these high outliers, we winsorized the false memories variable to the 95^th^ percentile. Winsorizing improved model fits but did not affect the statistical significance of our results.

Lastly, we quantified semantic similarity among the videos by using the Cluster Analysis function in *NVivo*. A.S. and G.M. produced detailed written descriptions of each video (available online: https://osf.io/xb7sq/), transcribing the narrative, setting, and character information. The similarity analysis filtered the written descriptions to exclude non-descriptive words (e.g., *the, and, in, on, under*), and then calculated pairwise Pearson correlations between videos on the basis of the frequency of the unique words used to describe each video. For each video, we calculated an overall semantic similarity score by averaging the correlation values; this metric summarized how much the content of a given video related to the rest of the stimulus set.

**Online Ratings of Stimulus Videos**

We recruited 3,913 participants online using Amazon’s Mechanical Turk. Participants were paid $0.50 to complete a 3-minute Qualtrics survey. Each participant was randomly assigned to view one stimulus video, first as the Full version and then as the Interrupted version. We included timing constraints to ensure that participants could not progress to the next page of the survey before the video had finished playing. Participants were excluded for the following reasons: (1) failing the attention check question (“If you are paying attention, choose 4 below.”), (2) failing the comprehension check question (“In general, not just in the video, is the emotion ‘happiness’ positive or negative?”), (3) video playback issues, or (4) prior exposure to the video. After exclusions, our sample size was 1,907 (20-41 raters per video). On 5-point Likert scales, participants rated how surprising each video felt when the ending was interrupted, as well as video memorability and emotional valence/intensity (Tables S14-S16). For each video, we averaged the surprise ratings from all Mechanical Turk participants to yield an average surprise score. These average per-video surprise scores (continuous variable) were then submitted to linear mixed effects regression models to predict memory outcomes in our laboratory sample (Tables S3).

**Exclusions**

In the Immediate group, two participants were excluded due to technical issues. In the Delayed group, three participants were excluded due to a counterbalancing error and audio playback problems, and two participants were excluded because they had previously completed a pilot version of the study. Additionally, one full run of fMRI data (14 trials) was excluded for one participant due to audio playback failure and excessive motion. On a trial-by-trial basis, videos were excluded if technical issues arose (e.g., audio issues) (10 trials), the participant was falling asleep (as determined by eyetracking) (20 trials), or the participant reported having seen the video prior to the experiment (103 trials). In total, there were 147 trials that were excluded for the above reasons (out of all 48 participants in both the Delayed and Immediate groups). The total number of excluded trials for Full and Interrupted videos was approximately equal (Full: 70; Interrupted: 77). Additionally, subsequently forgotten videos were excluded from single-trial brain-to-behavior analyses (63 trials across the 24 participants in the Delayed group). Overall, only 4.4% of all trials were excluded.

**Linear Mixed-Effects Regression**

All linear mixed-effects regression models reported in the main text included random intercepts for *subject* (identity of each participant) and *video* (identity of each stimulus item). In accordance with current best practices (1), we used the random effects structure that captured the maximal amount of complexity that was supported by the data (i.e., allowing model convergence and avoiding overfitting). Maximal models with all possible random slopes did not converge, so we incrementally simplified models by removing random slopes, then evaluated model fits by using a model comparison procedure (Likelihood Ratio Test with the Akaike Information Criterion). Full details about the random effects structure for each model are provided in the Supplementary Information, in each table description.

In R (v3.6), we constructed models with the *lme4* package (2) and evaluated significance with the *lmerTest* package (3). Variables for *reactivation type* and *group* were treated as factors, and all continuous variables were standardized/mean-centered. These model parameters applied to analysis of behavioral data, single-trial univariate neural activation, and temporal autocorrelation. Parameter estimates from all models are reported in the Supplementary Information Tables. All models converged successfully (BOBYQA controller, REML estimation, Satterthwaite degrees of freedom). Plots were generated with the packages *ggplot2* and *sjPlot* (4, 5). Simple slope estimates for each reactivation type condition (for significant interaction terms) were generated with the *emmeans* package (6).

**fMRI Preprocessing**

All data were preprocessed and analyzed using FSL v6.0, in conjunction with in-house R code (v3.6). Initial volumes were discarded by the scanner to allow for signal saturation. Preprocessing steps included fieldmap distortion correction, spatial realignment, removal of head-motion artifacts (six regressors), nuisance regression of average white matter and CSF timeseries, slice-timing correction for an interleaved multiband acquisition, and high-pass frequency filtering (120s). For native-space ROI analyses (single-trial univariate and autocorrelation analyses), data were minimally smoothed with a 2-mm kernel to preserve spatial specificity and multivariate information.

**Region of Interest Masks**

We used FreeSurfer (v6.0) (7) (<http://surfer.nmr.mgh.harvard.edu/>), to automatically create binarized hippocampal masks in each subject’s native space. Hippocampal masks were then manually inspected and segmented into ROIs for left anterior, left posterior, right anterior, and right posterior hippocampus. Anterior and posterior regions were split along the long-axis at the uncal apex. White matter masks were obtained with FSL segmentation utilities. Inferior Lateral Occipital Cortex (LOC) masks were taken from the Harvard-Oxford Cortical Atlas. VTA masks were taken from a probabilistic midbrain atlas developed by the Adcock lab (8). Basal forebrain masks were taken from the probabilistic cytoarchitectonic Julich-Brain atlas. We used ROIs for bilateral cholinergic nuclei Ch123, including the medial septal nucleus. This region (in contrast to Ch4) exhibits resting-state functional connectivity with the hippocampus (9). We investigated temporal signal-to-noise in the basal forebrain to ensure that results were not driven by noise (Supplementary Information, *Basal Forebrain SNR*). All standard space masks were transformed into native space for each functional run, using the inverse deformation field from preprocessing and registration.

*Table S1.* Parameter estimates from linear mixed effects regression models predicting (A) correct details and (B) false memories. Models included random intercepts for subjects and videos, random slopes for *reactivation type* for both subjects and videos, and random slopes for *group* for videos.

| **A) Dependent Variable: Correct Details** | | | |
| --- | --- | --- | --- |
| *Predictors* | *Estimates* | *CI* | *p* |
| (Intercept) | -0.01 | -0.16 – 0.14 | 0.891 |
| Reactivation Type | -0.07 ^**^ | -0.12 – -0.02 | **0.008** |
| Group | 0.16 ^*^ | 0.01 – 0.31 | **0.036** |
| Reactivation Type * Group | -0.01 | -0.04 – 0.02 | 0.495 |
| B) **Dependent Variable: False Memories** | | | |
| *Predictors* | *Estimates* | *CI* | *p* |
| (Intercept) | 0.01 | -0.09 – 0.11 | 0.842 |
| Reactivation Type | -0.05 ^**^ | -0.08 – -0.02 | **0.001** |
| Group | -0.36 ^***^ | -0.43 – -0.29 | **<0.001** |
| Reactivation Type * Group | 0.04 ^**^ | 0.01 – 0.07 | **0.009** |
| ** p<0.05   ** p<0.01   *** p<0.001* | | | |

**Trial Number Control**

To ensure that the effect of prediction error on memory updating was not driven by the first few trials (which are presumably most surprising), we also conducted a control analysis to test whether memory updating changed over the course of the experiment (effect of *trial number*). All parameter estimates are provided in Table S2. There was no main effect of trial number on false memories (β = 0.01, 95% CI [-0.03, 0.04], *t* = 0.48, *p* = .730), no interaction between trial number and reactivation type (β = 0.00, 95% CI [-0.03, 0.03], *t* = -0.09, *p* = .997), and no interaction among trial number, reactivation type, and group (β = 0.00, 95% CI [-0.03, 0.03], *t* = -0.40, *p* = .987). Overall, these null results suggest that the effect of Interrupted videos on false memories did not diminish over the course of the experiment.

*Table S2.* Parameter estimates from a linear mixed effects regression model predicting false memories from the variables *reactivation type* (Full vs. Interrupted), *group* (Delayed vs. Immediate), *trial number*, and all interactions. This analysis tested whether the effects of reactivation type on memory updating diminished over the course of the experiment (e.g., if interruptions become less surprising and therefore less effective). The model included random intercepts for subjects and videos, random slopes for *reactivation type* for both subjects and videos, and random slopes for *group* for videos.

| **Dependent Variable: False Memories** | | | |
| --- | --- | --- | --- |
| *Predictors* | *Estimates* | *CI* | *p* |
| (Intercept) | 0.01 | -0.09 – 0.11 | 0.841 |
| Reactivation Type | -0.05 ^**^ | -0.08 – -0.02 | **0.001** |
| Group | -0.36 ^***^ | -0.43 – -0.29 | **<0.001** |
| Trial | 0.01 | -0.02 – 0.04 | 0.631 |
| Reactivation Type * Group | 0.04 ^**^ | 0.01 – 0.07 | **0.009** |
| Reactivation Type * Trial | -0.00 | -0.03 – 0.03 | 0.927 |
| Group * Trial | -0.00 | -0.03 – 0.03 | 0.892 |
| Reactivation Type * Group * Trial | -0.01 | -0.04 – 0.02 | 0.693 |
| ** p<0.05   ** p<0.01   *** p<0.001* | | | |

**Confidence and Forgetting**

For each video, participants self-reported *confidence* ratings on a 5-point Likert scale (*not at all confident … very confident*). We used linear mixed effects regression to test whether confidence was influenced by group or reactivation type. Self-reported confidence ratings were slightly lower for interrupted videos, β = 0.03, 95% CI [0.001, 0.06], *t* = 2.13, *p* = .042. Neither the main effect of group (β = 0.11, 95% CI [-0.04, 0.25], *t* = 1.45, *p* = .155) nor the interaction term (β = -0.01, 95% CI [-0.04, 0.02], *t* = -0.57, *p* = .572) were significantly related to confidence ratings. Numerically, average confidence ratings were between “moderately confident” and “very confident” for all conditions (*M* = 3.76, *SD* = 0.85), suggesting that metamemory judgements were not very sensitive (Figure S1, left).

If a participant entirely failed to recall a video when cued with its name, the trial was scored as a *forgotten video*. We compared the average number of forgotten videos across conditions. Because there were few observations and this dependent measure is binary, we conducted a repeated-measures ANOVA on the sum of forgotten videos by-subject. There were more forgotten videos in the Delayed group than in the Immediate group, *F*_(1,46)_ = 10.36, *p* = .002, likely reflecting the additional 24-hour delay between reactivation and test in the Delayed group. Neither reactivation type (*F*_(1,46)_ = 2.77, *p* = .10) nor the interaction between group and reactivation type (*F*_(1,46)_ = 1.41, *p* = .24) were significantly related to forgotten videos. Thus, prediction error during memory reactivation did not significantly influence forgetting in either group (Figure S1, right).

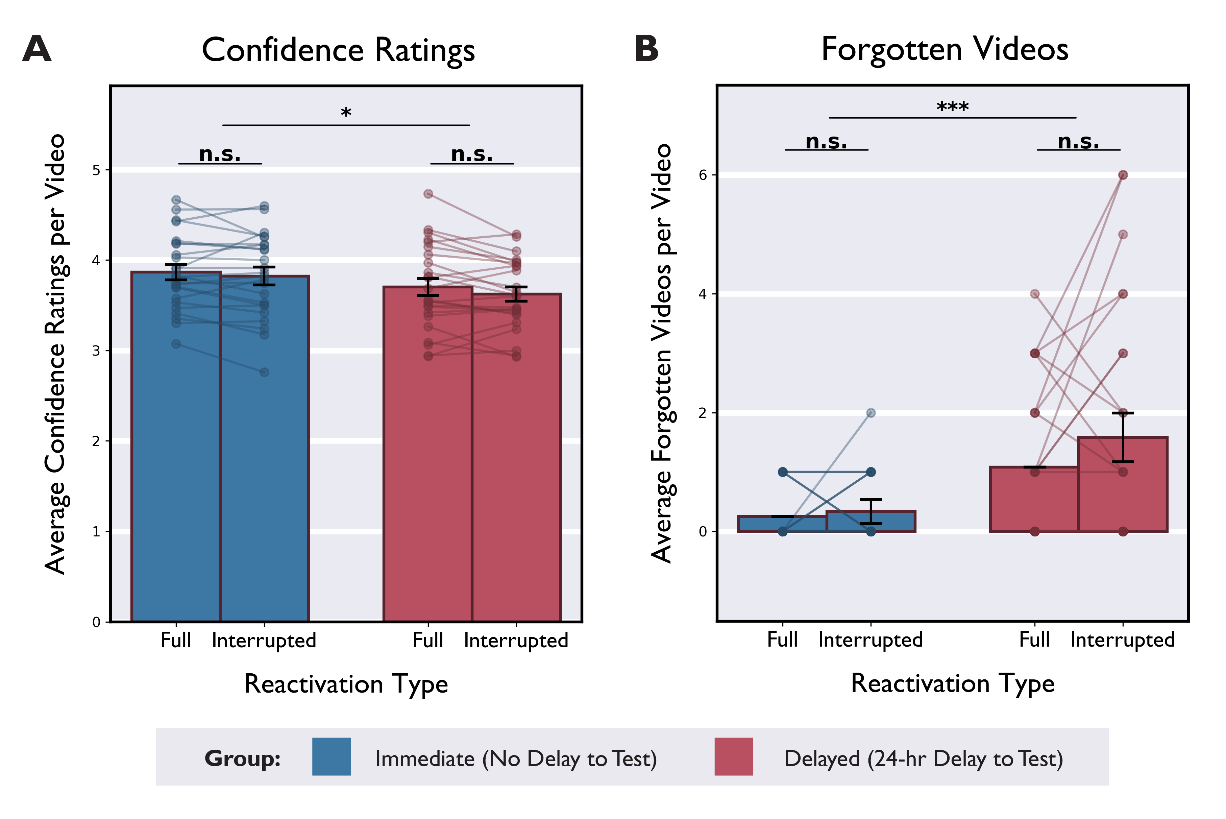

*Figure S1.* Average self-reported confidence ratings (left) and number of forgotten videos (right) subset by group (Delayed and Immediate) and reactivation type (Full and Interrupted). Dots indicate subject averages, and lines connect within-subjects measures. ** p<0.05   ** p<0.01   *** p<0.001*

*Table S3.* Parameter estimates from linear mixed effects regression models relating by-video surprise ratings (Supplementary Methods, *Online Ratings of Stimulus Videos*) to (A) false memories and (B) correct details. Models included random intercepts for subjects and videos, and random slopes for *reactivation type.*

| A) **Dependent Variable: False Memories** | | | |
| --- | --- | --- | --- |
| *Predictors* | *Estimates* | *CI* | *p* |
| (Intercept) | 0.01 | -0.09 – 0.11 | 0.842 |
| Reactivation Type | -0.05 ^**^ | -0.08 – -0.02 | **0.001** |
| Surprise | 0.07 | -0.00 – 0.15 | 0.068 |
| Group | -0.36 ^***^ | -0.43 – -0.29 | **<0.001** |
| Reactivation Type * Surprise | 0.00 | -0.03 – 0.03 | 0.843 |
| Reactivation Type * Group | 0.04 ^**^ | 0.01 – 0.07 | **0.009** |
| Surprise * Group | -0.03 ^*^ | -0.06 – -0.00 | **0.039** |
| Reactivation Type * Surprise * Group | 0.01 | -0.02 – 0.04 | 0.451 |
| B) **Dependent Variable: Correct Details** | | | |
| *Predictors* | *Estimates* | *CI* | *p* |
| (Intercept) | -0.01 | -0.16 – 0.14 | 0.895 |
| Reactivation Type | -0.07 ^**^ | -0.12 – -0.02 | **0.008** |
| Surprise | 0.03 | -0.01 – 0.07 | 0.199 |
| Group | 0.16 ^*^ | 0.02 – 0.31 | **0.036** |
| Reactivation Type * Surprise | -0.00 | -0.05 – 0.04 | 0.841 |
| Reactivation Type * Group | -0.01 | -0.04 – 0.02 | 0.490 |
| Surprise * Group | -0.01 | -0.04 – 0.02 | 0.509 |
| Reactivation Type * Surprise * Group | 0.00 | -0.03 – 0.03 | 0.786 |
| ** p<0.05   ** p<0.01   *** p<0.001* | | | |

*Table S4.* Parameter estimates from linear mixed effects regression models relating by-video semantic similarity scores to (A) false memories and (B) correct details. Models included random intercepts for subjects and videos, and random slopes for *reactivation type.*

| A) **Dependent Variable: False Memories** | | | |
| --- | --- | --- | --- |
| *Predictors* | *Estimates* | *CI* | *p* |
| (Intercept) | 0.01 | -0.09 – 0.10 | 0.855 |
| Reactivation Type | -0.05 ^***^ | -0.08 – -0.02 | **0.001** |
| Similarity | 0.11 ^**^ | 0.04 – 0.19 | **0.003** |
| Group | -0.36 ^***^ | -0.43 – -0.29 | **<0.001** |
| Reactivation Type * Similarity | -0.03 ^*^ | -0.06 – -0.00 | **0.038** |
| Reactivation Type * Group | 0.04 ^**^ | 0.01 – 0.07 | **0.010** |
| Similarity * Group | -0.04 ^**^ | -0.07 – -0.01 | **0.006** |
| Reactivation Type * Similarity * Group | 0.02 | -0.02 – 0.05 | 0.323 |

| C) **Dependent Variable: Correct Details** | | | |
| --- | --- | --- | --- |
| *Predictors* | *Estimates* | *CI* | *p* |
| (Intercept) | -0.01 | -0.16 – 0.14 | 0.894 |
| Reactivation Type | -0.07 ^**^ | -0.12 – -0.02 | **0.008** |
| Similarity | -0.05 ^*^ | -0.09 – -0.01 | **0.025** |
| Group | 0.16 ^*^ | 0.02 – 0.31 | **0.035** |
| Reactivation Type * Similarity | 0.01 | -0.04 – 0.05 | 0.777 |
| Reactivation Type * Group | -0.01 | -0.04 – 0.02 | 0.482 |
| Similarity * Group | -0.00 | -0.03 – 0.03 | 0.835 |
| Reactivation Type * Similarity * Group | 0.01 | -0.02 – 0.04 | 0.542 |
| ** p<0.05   ** p<0.01   *** p<0.001* | | | |

**Behavioral Variance Control**

Overall, participants in the Immediate group reported fewer false memories (average of 0.43 false memories per video, relative to 1.14 in the Delayed group) and showed lower variance for this measure (standard deviation of 0.09, relative to 0.17 in the Delayed group). One possibility is that there was insufficient variance to detect any effect of reactivation type in this group. In other words, there may be a floor effect that would make it difficult to detect condition differences that may not depend on a delay-to-test. To investigate this possibility, we conducted a control analysis that excluded low-variance subjects from the Immediate group.

First, for each subject we calculated the standard deviation for false memories across all trials. Rank-ordering these variance scores revealed that five subjects in the Immediate group had very low standard deviations (as well as fairly few false memories in total). Second, we explored whether it was possible to detect *any* condition differences in the Immediate group. As reported in the Results and Table S1, we observed that for participants in the Delayed group, there was a consistent bias towards more false memories for Interrupted > Full videos (21/24 participants showed this directional bias). Although some participants in the Immediate group reported more false memories for Interrupted > Full videos (14/24), others showed the opposite trend (7/24) or no difference (3/24).

To investigate whether our paradigm produced enough variance to detect *any* condition differences in false memories (in either direction), we calculated the absolute value of these difference scores. Focusing on the absolute value difference between conditions, rather than the total number of false memories, may offer insight into whether our paradigm is sensitive enough to detect any possible condition differences. If a participant reported few false memories and showed low variance across trials, we would expect no difference between conditions.

We found that the same five participants in the Immediate group who showed very low variance in the false memories measure also showed the smallest (absolute value) differences in false memories between Full and Interrupted conditions. Overall, we observed that these five participants had few false memories and low variance in their responses, which may have affected group-level statistics as well.

Therefore, we conducted a control analysis in which we excluded these five low-variance participants from the Immediate group. As a result, we were able to compare the effects of Group and Reactivation Type *among the subset of participants who showed any difference in false memories* between Reactivation Type conditions.

We found that omitting these low-variance subjects from the Immediate group did not appreciably change our results. As reported in the main text (Results, “*Prediction Error Increased False Memories*”), we found a significant interaction between Group and Reactivation Type (β = 0.04, 95% CI [0.01, 0.07], *t* = 2.25, *p* = .025), such that Delayed group participants reported more false memories for Interrupted videos than Full videos (β = -0.09, *t* = -3.50, *p* < .001), but Immediate group participants showed no effect of reactivation type (β = -0.01, *t* = -0.74, *p* = .461). There was also a significant main effect of group (Delayed > Immediate; β = -0.35, 95% CI [-0.43, -0.27], *t* = -8.55, *p* < .001) and a significant main effect of type (Interrupted > Full; β = -0.06, 95% CI [-0.09, -0.02], *t* = -3.17, *p* = .002), both driven by the Delayed group.

**Whole-Brain Analysis**

Functional and anatomical data were aligned and spatially normalized to MNI space. Preprocessing steps are described in the Methods section of the main text. The normalized functional data were spatially smoothed using a 3D 8-mm full-width-half-maximum (FWHM) Gaussian kernel. BOLD activation for each condition was averaged within-run for each subject, across runs within-subjects, and finally across all subjects. Cluster inference was conducted with an initial cluster-forming threshold of *p* < .001 and a final family-wise error corrected significance threshold of *p* < .05, derived from Gaussian Random Field Theory. Whole-brain activation maps depict *z*-stat values after correction for multiple comparisons (Figure S5). Group-level analyses were conducted with FSL’s FLAME1 mixed-effects function, with automatic outlier deweighting.

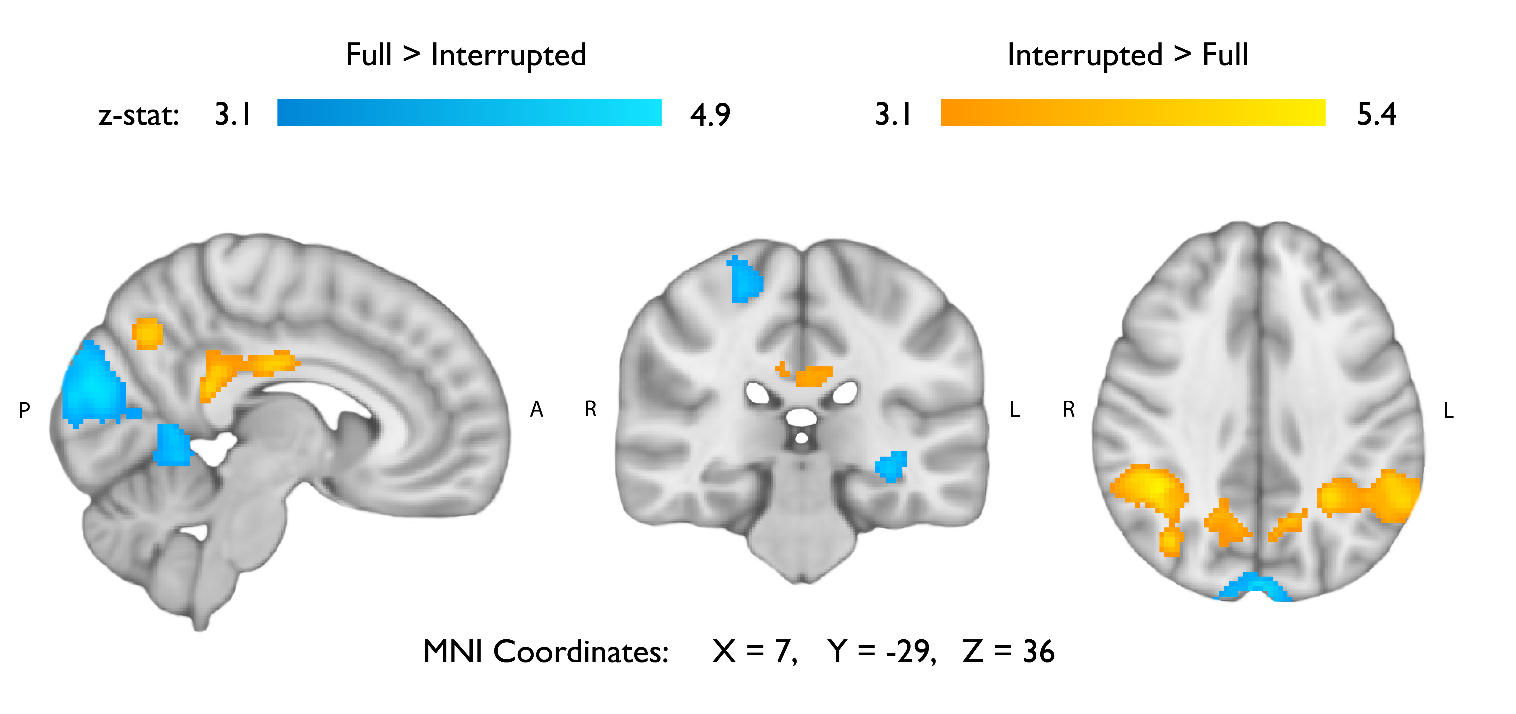
We contrasted Full and Interrupted trials to obtain a whole-brain map of averaged group-level BOLD activation (Figure S2, Table S5). We found distributed clusters of neural activation that differed between the conditions, supporting the idea that prediction error influences post-event processing. A large area of the lateral parietal cortex demonstrated significantly greater activation after interrupted videos than after full-length videos. These bilateral parietal clusters included portions of the angular gyrus, superior parietal lobule, and supramarginal gyrus. Other regions that were significantly activated more after Interrupted videos than Full videos included the precuneus and posterior-mid cingulate cortex. Clusters in the left hippocampus, cerebellum, and occipital cortex were significantly activated more after Full videos than Interrupted videos. A table reporting significant clusters is provided in Table S5.

*Figure S2.* Whole-brain univariate activation results. Contrasts compare average BOLD activation during the post-event fixation period following Interrupted and Full-Length videos.

*Table S5.* Statistics and coordinates of significant clusters for whole-brain contrasts of the fixation period following Full and Interrupted videos. Statistical thresholding was set at *z* = 3.1 (*p* < .001) for initial cluster formation, and *p* < .05 for subsequent family-wise error correction of clusters.

| **Contrast** | **Regions** | **N Voxels** | **P-Value** | **Max Z-Stat** | **Peak X** | **Peak Y** | **Peak Z** | **Mean COPE** |
| --- | --- | --- | --- | --- | --- | --- | --- | --- |
| Full > Interrupted | Left Hippocampus, Bilateral Supracalcarine Sulcus, Bilateral Cuneal Cortex, Bilateral Occipital Cortex, Bilateral Lingual Gyrus | 5470 | 1.20E-17 | 4.94 | 2 | -82 | 16 | 25.1 |
| Full > Interrupted | Right Postcentral Gyrus | 251 | 0.0438 | 4.3 | 28 | -34 | 76 | 15.5 |
| Interrupted > Full | Right Superior Lateral Occipital Cortex, Right Superior Parietal Lobule, Right Angular Gyrus, Right Supramarginal Gyrus | 2645 | 8.66E-11 | 5.4 | 42 | -56 | 60 | 30 |
| Interrupted > Full | Left Superior Lateral Occipital Cortex, Left Superior Parietal Lobule, Left Angular Gyrus, Left Supramarginal Gyrus | 1748 | 5.96E-08 | 4.85 | -36 | -62 | 60 | 26.6 |
| Interrupted > Full | Posterior Cingulate Cortex, Mid Cingulate Cortex | 624 | 0.000548 | 4.82 | 2 | -24 | 28 | 22 |
| Interrupted > Full | Bilateral Precuneus | 481 | 0.00256 | 4.54 | 0 | -70 | 44 | 23.4 |
| Interrupted > Full | Right Cerebellum | 380 | 0.00832 | 4.48 | 30 | -64 | -30 | 18.4 |
| Interrupted > Full | Left Cerebellum | 363 | 0.0102 | 4.94 | -30 | -60 | -30 | 16.5 |

*Table S6.* Parameter estimates from a linear mixed effects regression model predicting trial-wise univariate hippocampal activation in four non-overlapping ROIs (left anterior, right anterior, left posterior, and right posterior hippocampus). Fixed effects were *reactivation type* (Full vs. Interrupted), *false memories, hemisphere* (left vs. right), *axis* (anterior vs. posterior), and all interactions. Model included random intercepts for subjects and videos; random slopes for *reactivation type*, *false memories*, *hemisphere*, and *axis* for each subject; and random slopes for *reactivation type* for each video. Boldface indicates statistically significant parameters.

| **Dependent Variable: Hippocampal Activation** | | | |
| --- | --- | --- | --- |
| *Predictors* | *Estimates* | *CI* | *p* |
| (Intercept) | -0.01 | -0.10 – 0.08 | 0.873 |
| Reactivation Type | 0.04 | -0.00 – 0.09 | 0.069 |
| False Memories | -0.01 | -0.05 – 0.03 | 0.640 |
| Hemisphere | 0.00 | -0.02 – 0.03 | 0.858 |
| Axis | 0.04 ^**^ | 0.01 – 0.06 | **0.002** |
| Reactivation Type * False Memories | -0.06 ^***^ | -0.09 – -0.03 | **<0.001** |
| Reactivation Type * Hemisphere | 0.01 | -0.02 – 0.03 | 0.640 |
| Reactivation Type * Axis | -0.00 | -0.03 – 0.02 | 0.830 |
| False Memories * Hemisphere | -0.02 | -0.04 – 0.01 | 0.184 |
| False Memories * Axis | 0.01 | -0.01 – 0.03 | 0.432 |
| Reactivation Type * False Memories * Hemisphere | 0.01 | -0.02 – 0.03 | 0.497 |
| Reactivation Type * False Memories * Axis | -0.01 | -0.03 – 0.02 | 0.642 |
| ** p<0.05   ** p<0.01   *** p<0.001* | | | |

*Table S7.* Parameter estimates from a linear mixed effects regression model predicting trial-wise univariate hippocampal activation. Fixed effects were *reactivation type* (Full vs. Interrupted), *false memories, correct details, hemisphere* (left vs. right), *axis* (anterior vs. posterior), and relevant interactions. Model included random intercepts for subjects and videos; random slopes for *reactivation type*, *false memories*, and *correct details* for each subject; and random slopes for *reactivation type* for each video. Boldface indicates statistically significant parameters.

| **Dependent Variable: Hippocampal Activation** | | | |
| --- | --- | --- | --- |
| *Predictors* | *Estimates* | *CI* | *p* |
| (Intercept) | -0.00 | -0.10 – 0.10 | 0.997 |
| Reactivation Type | 0.04 | -0.00 – 0.09 | 0.074 |
| False Memories | -0.01 | -0.04 – 0.03 | 0.783 |
| Correct Details | -0.02 | -0.08 – 0.03 | 0.474 |
| Hemisphere | 0.00 | -0.02 – 0.03 | 0.766 |
| Axis | 0.04 ^**^ | 0.02 – 0.06 | **0.001** |
| Reactivation Type * False Memories | -0.06 ^***^ | -0.08 – -0.03 | **<0.001** |
| Reactivation Type * Correct Details | 0.03 ^*^ | 0.00 – 0.06 | **0.036** |
| False Memories * Correct Details | -0.01 | -0.04 – 0.02 | 0.419 |
| Reactivation Type * False Memories * Correct Details | 0.04 ^*^ | 0.01 – 0.06 | **0.013** |
| ** p<0.05   ** p<0.01   *** p<0.001* | | | |

*Table S8.* Parameter estimates from a linear mixed effects regression model predicting trial-wise univariate hippocampal activation, with the addition of basal forebrain moderation. Fixed effects were *reactivation type* (Full vs. Interrupted), *false memories, basal forebrain activation (BF), hemisphere* (left vs. right), *axis* (anterior vs. posterior), and relevant interactions. Model included random intercepts for subjects and videos; random slopes for *reactivation type*, *false memories*, and *basal forebrain* for each subject; and random slopes for *reactivation type* for each video. Boldface indicates statistically significant parameters.

| **Dependent Variable: Hippocampal Activation** | | | |
| --- | --- | --- | --- |
| *Predictors* | *Estimates* | *CI* | *p* |
| (Intercept) | -0.01 | -0.10 – 0.08 | 0.815 |
| Reactivation Type | 0.04 | -0.01 – 0.08 | 0.123 |
| False Memories | -0.01 | -0.04 – 0.03 | 0.718 |
| BF | 0.10 ^*^ | 0.02 – 0.19 | **0.029** |
| Hemisphere | 0.00 | -0.02 – 0.03 | 0.892 |
| Axis | 0.04 ^**^ | 0.02 – 0.06 | **0.001** |
| Reactivation Type * False Memories | -0.06 ^***^ | -0.09 – -0.03 | **<0.001** |
| Reactivation Type * BF | -0.02 | -0.05 – 0.00 | 0.068 |
| False Memories * BF | -0.02 | -0.05 – 0.01 | 0.132 |
| Reactivation Type * False Memories * BF | -0.04 ^**^ | -0.06 – -0.01 | **0.008** |
| Reactivation Type * False Memories * BF * Hemisphere | 0.01 | -0.02 – 0.03 | 0.654 |
| Reactivation Type * False Memories * BF * Axis | 0.01 | -0.01 – 0.04 | 0.367 |
| ** p<0.05   ** p<0.01   *** p<0.001* | | | |

*Table S9.* Parameter estimates from a linear mixed effects regression model predicting trial-wise univariate hippocampal activation. Fixed effects were *reactivation type* (Full vs. Interrupted), *false memories, VTA activation, hemisphere* (left vs. right), *axis* (anterior vs. posterior), and relevant interactions. Model included random intercepts for subjects and videos; random slopes for *reactivation type*, *false memories*, and *VTA* for each subject; and random slopes for *reactivation type* for each video. Boldface indicates statistically significant parameters.

| **Dependent Variable: Hippocampal Activation** | | | |
| --- | --- | --- | --- |
| *Predictors* | *Estimates* | *CI* | *p* |
| (Intercept) | -0.01 | -0.09 – 0.07 | 0.863 |
| Reactivation Type | 0.05 ^*^ | 0.00 – 0.10 | **0.039** |
| False Memories | -0.01 | -0.05 – 0.03 | 0.607 |
| VTA | 0.15 ^**^ | 0.05 – 0.25 | **0.006** |
| Hemisphere | 0.00 | -0.02 – 0.03 | 0.878 |
| Axis | 0.04 ^**^ | 0.02 – 0.06 | **0.001** |
| Reactivation Type * False Memories | -0.06 ^***^ | -0.08 – -0.03 | **<0.001** |
| Reactivation Type * VTA | 0.01 | -0.02 – 0.03 | 0.678 |
| False Memories * VTA | 0.02 | -0.01 – 0.05 | 0.141 |
| Reactivation Type * False Memories * VTA | -0.00 | -0.03 – 0.02 | 0.747 |
| Reactivation Type * False Memories * VTA * Hemisphere | 0.01 | -0.02 – 0.03 | 0.465 |
| Reactivation Type * False Memories * VTA * Axis | -0.01 | -0.04 – 0.01 | 0.414 |
| ** p<0.05   ** p<0.01   *** p<0.001* | | | |

*Table S10.* Parameter estimates from linear mixed effects regression models predicting trial-wise univariate basal forebrain activation (A) and VTA activation (B). Fixed effects were *reactivation type* (Full vs. Interrupted), *false memories,* and the interaction. Models included random intercepts for subjects and videos; random slopes for *reactivation type* and *false memories* for each subject; and random slopes for *reactivation type* for each video. Boldface indicates statistically significant parameters.

| A) **Dependent Variable: Basal Forebrain Activation** | | | |
| --- | --- | --- | --- |
| *Predictors* | *Estimates* | *CI* | *p* |
| (Intercept) | -0.01 | -0.14 – 0.13 | 0.923 |
| Reactivation Type | 0.02 | -0.04 – 0.07 | 0.559 |
| False Memories | -0.01 | -0.06 – 0.05 | 0.822 |
| Reactivation Type * False Memories | 0.00 | -0.05 – 0.05 | 0.955 |
| B) **Dependent Variable: VTA Activation** | | | |
| *Predictors* | *Estimates* | *CI* | *p* |
| (Intercept) | -0.02 | -0.16 – 0.13 | 0.829 |
| Reactivation Type | -0.03 | -0.08 – 0.02 | 0.260 |
| False Memories | 0.01 | -0.04 – 0.06 | 0.777 |
| Reactivation Type * False Memories | -0.02 | -0.07 – 0.03 | 0.403 |
| ** p<0.05   ** p<0.01   *** p<0.001* | | | |

*Table S11.* Parameter estimates from linear mixed effects regression models examining hippocampal autocorrelation. A) Predicting hippocampal autocorrelation over the course of video playback. Segments indicate 5-second bins during the video stimulus. Model included random intercepts for subjects and videos; random slopes for segment, hemisphere, and axis for each subject. B) Predicting average Post-Video change in autocorrelation from reactivation type (subtracting average autocorrelation from the 5s pre-offset from average autocorrelation from the 5s bin post-offset). Model included random intercepts for subjects and videos, and random slopes for reactivation type.

| A) **Dependent Variable: Hippocampal Autocorrelation** | | | |
| --- | --- | --- | --- |
| *Predictors* | *Estimates* | *CI* | *p* |
| (Intercept) | 0.01 | -0.21 – 0.22 | 0.955 |
| Video Segment | 0.025 ^**^ | 0.01 – 0.04 | **0.002** |
| Hemisphere | -0.00 | -0.03 – 0.02 | 0.750 |
| Axis | 0.03 | -0.01 – 0.07 | 0.184 |
| Video Segment * Hemisphere | 0.00 | -0.01 – 0.01 | 0.365 |
| Video Segment * Axis | 0.01 | -0.00 – 0.02 | 0.244 |
| Hemisphere * Axis | -0.01 | -0.02 – 0.00 | 0.105 |
| Segment * Hemisphere * Axis | -0.00 | -0.01 – 0.01 | 0.805 |

| B) **Dependent Variable: Post-Video Change in Hippocampal Autocorrelation** | | | |
| --- | --- | --- | --- |
| *Predictors* | *Estimates* | *CI* | *p* |
| (Intercept) | 0.02 | -0.03 – 0.07 | 0.421 |
| Reactivation Type | 0.04 ^*^ | 0.01 – 0.07 | **0.038** |
| Hemisphere | -0.00 | -0.03 – 0.02 | 0.826 |
| Axis | -0.02 | -0.05 – 0.00 | 0.058 |
| Reactivation Type * Hemisphere | 0.00 | -0.02 – 0.03 | 0.723 |
| Reactivation Type * Axis | -0.00 | -0.03 – 0.02 | 0.901 |
| ** p<0.05   ** p<0.01   *** p<0.001* | | | |

*Table S12*. Parameter estimates from linear mixed effects regression models relating hippocampal autocorrelation to subsequent false memories. A) Predicting subsequent false memories from reactivation type (Full vs. Interrupted), univariate hippocampal activation (HPC), post-video change in hippocampal autocorrelation (Autocor Change), hemisphere, axis, and interaction terms. Model included random intercepts for subjects and videos; random slopes for reactivation type, univariate hippocampal activation, and autocor change for each subject; and random slopes for reactivation type for each video. B) Expanded model that adds parameters for univariate basal forebrain activation, and accompanying random slopes for basal forebrain activation for each subject. C) Expanded model that adds parameters for univariate VTA activation, and accompanying random slopes for VTA activation for each subject. Boldface indicates statistically significant parameters.

| A) **Dependent Variable: False Memories** | | | |
| --- | --- | --- | --- |
| *Predictors* | *Estimates* | *CI* | *p* |
| (Intercept) | 0.03 | -0.11 – 0.18 | 0.653 |
| Reactivation Type | -0.10 ^***^ | -0.15 – -0.04 | **0.001** |
| HPC | -0.00 | -0.04 – 0.03 | 0.917 |
| Autocor Change | -0.02 | -0.05 – 0.01 | 0.247 |
| Hemisphere | -0.00 | -0.02 – 0.02 | 0.995 |
| Axis | 0.00 | -0.02 – 0.03 | 0.799 |
| Autocor Change * Hemisphere | 0.01 | -0.02 – 0.03 | 0.517 |
| Autocor Change * Axis | -0.00 | -0.02 – 0.03 | 0.942 |
| Reactivation Type * HPC | -0.05 ^***^ | -0.07 – -0.02 | **<0.001** |
| Reactivation Type * Autocor Change | 0.04 ^**^ | 0.02 – 0.07 | **0.001** |
| Reactivation Type * Hemisphere | -0.00 | -0.02 – 0.02 | 0.895 |
| Reactivation Type * Axis | 0.00 | -0.02 – 0.03 | 0.778 |
| Reactivation Type * Autocor Change * Hemisphere | -0.02 | -0.05 – 0.00 | 0.074 |
| Reactivation Type * Autocor Change * Axis | -0.02 | -0.04 – 0.01 | 0.128 |

| B) **Dependent Variable: False Memories** | | | |
| --- | --- | --- | --- |
| *Predictors* | *Estimates* | *CI* | *p* |
| (Intercept) | 0.04 | -0.10 – 0.19 | 0.580 |
| Reactivation Type | -0.09 ^***^ | -0.13 – -0.06 | **<0.001** |
| BF | -0.01 | -0.06 – 0.04 | 0.768 |
| HPC | -0.00 | -0.04 – 0.04 | 0.974 |
| Autocor Change | -0.02 | -0.05 – 0.01 | 0.209 |
| Hemisphere | 0.00 | -0.02 – 0.02 | 0.983 |
| Axis | 0.00 | -0.02 – 0.03 | 0.818 |
| Reactivation Type * BF | -0.02 | -0.04 – 0.01 | 0.127 |
| Autocor Change * Hemisphere | 0.01 | -0.02 – 0.03 | 0.439 |
| Autocor Change * Axis | 0.00 | -0.02 – 0.03 | 0.877 |
| Reactivation Type * HPC | -0.05 ^***^ | -0.07 – -0.02 | **<0.001** |
| Reactivation Type * Autocor Change | 0.04 ^***^ | 0.02 – 0.07 | **0.001** |
| Reactivation Type * Hemisphere | -0.00 | -0.02 – 0.02 | 0.979 |
| Reactivation Type * Axis | 0.00 | -0.02 – 0.03 | 0.723 |
| BF * HPC | -0.01 | -0.03 – 0.01 | 0.512 |
| BF * Autocor Change | 0.02 | -0.00 – 0.05 | 0.078 |
| BF * Hemisphere | 0.00 | -0.02 – 0.02 | 0.986 |
| BF * Axis | 0.00 | -0.02 – 0.02 | 0.871 |
| Reactivation Type * Autocor Change * Hemisphere | -0.03 ^*^ | -0.05 – -0.00 | **0.046** |
| Reactivation Type * Autocor Change * Axis | -0.02 | -0.05 – 0.00 | 0.115 |
| BF * Autocor Change * Hemisphere | -0.01 | -0.03 – 0.02 | 0.575 |
| BF * Autocor Change * Axis | -0.01 | -0.04 – 0.01 | 0.243 |
| Reactivation Type * HPC * BF | -0.03 ^*^ | -0.05 – -0.01 | **0.011** |
| Reactivation Type * Autocor Change * BF | 0.03 ^*^ | 0.01 – 0.06 | **0.011** |
| Reactivation Type * BF * Hemisphere | -0.00 | -0.02 – 0.02 | 0.885 |
| Reactivation Type * BF * Axis | 0.01 | -0.02 – 0.03 | 0.645 |
| Reactivation Type * Autocor Change * BF * Hemisphere | -0.01 | -0.03 – 0.02 | 0.653 |
| Reactivation Type * Autocor Change * BF * Axis | 0.01 | -0.02 – 0.03 | 0.665 |
| C) **Dependent Variable: False Memories** | | | |
| *Predictors* | *Estimates* | *CI* | *p* |
| (Intercept) | 0.04 | -0.11 – 0.19 | 0.622 |
| Reactivation Type | -0.11 ^**^ | -0.17 – -0.05 | **0.001** |
| VTA | -0.00 | -0.06 – 0.06 | 0.978 |
| HPC | -0.00 | -0.04 – 0.03 | 0.805 |
| Autocor Change | -0.02 | -0.06 – 0.01 | 0.228 |
| Hemisphere | -0.00 | -0.02 – 0.02 | 0.945 |
| Axis | 0.00 | -0.02 – 0.03 | 0.776 |
| Reactivation Type * VTA | -0.01 | -0.03 – 0.02 | 0.601 |
| Autocor Change * Hemisphere | 0.01 | -0.01 – 0.04 | 0.363 |
| Autocor Change * Axis | 0.00 | -0.02 – 0.03 | 0.748 |
| Reactivation Type * HPC | -0.05 ^***^ | -0.07 – -0.02 | **<0.001** |
| Reactivation Type * Autocor Change | 0.04 ^**^ | 0.01 – 0.06 | **0.002** |
| Reactivation Type * Hemisphere | -0.00 | -0.02 – 0.02 | 0.840 |
| Reactivation Type * Axis | 0.00 | -0.02 – 0.03 | 0.781 |
| VTA * HPC | 0.00 | -0.02 – 0.02 | 0.799 |
| VTA * Autocor Change | -0.02 | -0.04 – 0.01 | 0.187 |
| VTA * Hemisphere | -0.00 | -0.02 – 0.02 | 0.937 |
| VTA * Axis | -0.00 | -0.03 – 0.02 | 0.791 |
| Reactivation Type * Autocor Change * Hemisphere | -0.03 ^*^ | -0.05 – -0.00 | **0.043** |
| Reactivation Type * Autocor Change * Axis | -0.02 | -0.05 – 0.00 | 0.073 |
| VTA * Autocor Change * Hemisphere | 0.00 | -0.02 – 0.03 | 0.702 |
| VTA * Autocor Change * Axis | 0.01 | -0.01 – 0.04 | 0.192 |
| Reactivation Type * VTA * HPC | -0.02 ^*^ | -0.04 – -0.00 | **0.048** |
| Reactivation Type * VTA * Autocor Change | -0.01 | -0.03 – 0.02 | 0.587 |
| Reactivation Type * VTA * Hemisphere | 0.00 | -0.02 – 0.02 | 0.893 |
| Reactivation Type * VTA * Axis | -0.00 | -0.02 – 0.02 | 0.971 |
| Reactivation Type * VTA * Autocor Change * Hemisphere | -0.00 | -0.02 – 0.02 | 0.826 |
| Reactivation Type * VTA * Autocor Change * Axis | -0.01 | -0.03 – 0.01 | 0.434 |
| ** p<0.05   ** p<0.01   *** p<0.001* | | | |

*Table S13.* Names and brief descriptions (hints provided if necessary) for each stimulus video.

| **Video Name** | **Description/Hint** |
| --- | --- |
| Archery | Somebody is trying to shoot somebody with a bow and arrow, in a forest. |
| Baking Soda Volcano | A family is demonstrating how to make a baking soda volcano erupt. |
| Balloon Box | Somebody receives a Valentine's Day gift. |
| Baseball | Some people were playing baseball, and the batter hits a homerun. |
| Basketball | A basketball player does a slam dunk. |
| Biking | There are people on bicycles racing. |
| Birthday Dinner | Somebody has a birthday dinner with family, then a celebration in the living room. |
| Bumblebee | Somebody is getting arrested, but the person is afraid of a bumblebee. |
| Breakfast Plate | Somebody makes another person breakfast, but they do not like the breakfast. |
| Breaking Dishes | Somebody breaks some dishes in a cabinet. |
| Bridge Collapsing | A bridge cracks and falls apart. |
| Bridge Jump | Some people in a car try to drive and jump over a broken bridge. |
| Cannon | Some people fire a cannon during a battle. |
| Canoe Over Waterfall | Some people in a canoe go over a waterfall. |
| Car Crash | Somebody driving a car stops at an intersection and gets hit by another car. |
| Car Explosion | Somebody shoots at a car and it explodes. |
| Cliff Jump | Somebody jumps off a cliff and uses a rope to swing. |
| Construction Site | Somebody falls through the floor in a construction site. |
| Cruise Ship Crash | A cruise ship crashes into another boat. |
| Defusing Bomb | Some people are trying to defuse a bomb as the timer is counting down. |
| Diving | This is a diving competition at a swimming pool. |
| Drop Tower | Some people ride a drop tower at an amusement park. |
| Dunk Tank | Somebody throws a ball to dunk another person in a dunk tank. |
| Earthquake | An earthquake shakes a building and makes it fall. |
| Fire Alarm Sprinklers | Somebody sets off the sprinklers in the ceiling by using a lighter. |
| Fire Escape Breaking | Some people are on a fire escape outside an apartment building. |
| Fire Experiment | Some people are trying to make a fire really big. |
| Firefighter Ladder | Some firefighters are putting out a fire, but the ladder falls. |
| Food Fight | Somebody starts a food fight in a cafeteria. |
| Gambling | Somebody is gambling in a casino, rolling dice. |
| Graduation | Some people are having a graduation ceremony. |
| Horse Jumping | A horse and rider are in a competition where they jump over obstacles. |
| Hostages | Some people with guns are trying to save some hostages in a building. |
| Hunting | Somebody is hunting an animal. |
| Ice Breaking | Somebody is jumping on a frozen pool. |
| Jousting | Some people on horses are in a jousting tournament, with spears. |
| Kitchen Fire | Somebody is demonstrating what to do when there is a kitchen fire. |
| Lightning | There are some lightning sparks indoors, and then a big lightning strike outside. |
| Magic | A magician is doing a trick about changing the dollar value of money. |
| Minefield | Somebody is running across a minefield, trying not to step on the wires. |
| Motorized Chair | Somebody steals a motorized chair from somebody else. |
| Olympic Torch | Somebody uses a torch to light the flame at the Olympics |
| Olympic Vault | Somebody vaults over a hurdle and does some flips. |
| Orchestra Concert | An orchestra performs in a concert hall. |
| Plane Crash | A pilot and co-pilot crash a plane. |
| Pole Vault | An athlete uses a pole to jump over a tall bar. |
| Proposal | A woman proposes to a man. |
| Punch | Somebody commands people to punch another person. |
| Racecars | Some cars are racing on a road. |
| Raining Money | A magician does a trick to make it rain money. |
| Rescue Dog | A dog saves somebody who is drowning |
| Rollercoaster | A rollercoaster stops upside down. |
| Rooftop Chase | Somebody is chasing somebody else across rooftops. |
| Running Hug | Two people stare at each other and then run and hug. |
| Running Race | Athletes are running a race on a track. |
| Shot Put | An athlete throws a metal ball in a competition. |
| Shooting Range | Somebody shoots at a target that looks like the outline of a person. |
| Skateboard | Somebody rides a skateboard down the street. |
| Skydiving | Somebody skydives out of an airplane, but needs a push. |
| Slap | Somebody asks somebody else to slap them. |
| Sniper | A sniper is trying to shoot somebody. |
| Soccer | Two teams are playing soccer, and the goalie throws the ball. |
| Space Shuttle | A rocket is taking off. |
| Surprise Party | Somebody comes home and other people are waiting for a surprise party. |
| Swimming Pool Push | Somebody pushes somebody else into a swimming pool. |
| Train | Somebody is trying to jump onto a moving train. |
| Underwater Window | An underwater window breaks and floods a room. |
| Volcano Erupting | A family is trying to drive away while a volcano is erupting. |
| Wall Flip | Somebody is learning how to do a flip by kicking off the wall. |
| Wedding | There is a wedding and the audience members are talking. |

*Table S14.* Results from linear mixed effects regression models predicting memory outcomes from emotional valence ratings, reactivation type, and group. The model included random intercepts for subjects and videos, random slopes for *reactivation type* for both subjects and videos, and random slopes for *group* for videos.

| A) **Dependent Variable: Correct Details** | | | |
| --- | --- | --- | --- |
| *Predictors* | *Estimates* | *CI* | *p* |
| (Intercept) | -0.01 | -0.16 – 0.14 | 0.893 |
| Reactivation Type | -0.07 ^**^ | -0.12 – -0.02 | **0.008** |
| Valence | 0.01 | -0.03 – 0.05 | 0.660 |
| Group | 0.16 ^*^ | 0.02 – 0.31 | **0.036** |
| Reactivation Type * Valence | 0.01 | -0.04 – 0.06 | 0.673 |
| Reactivation Type * Group | -0.01 | -0.04 – 0.02 | 0.485 |
| Valence * Group | -0.02 | -0.05 – 0.01 | 0.191 |
| Reactivation Type * Valence * Group | 0.01 | -0.02 – 0.03 | 0.712 |
| **B) Dependent Variable: False Memories** | | | |
| *Predictors* | *Estimates* | *CI* | *p* |
| (Intercept) | 0.01 | -0.09 – 0.11 | 0.842 |
| Reactivation Type | -0.05 ^**^ | -0.08 – -0.02 | **0.001** |
| Valence | 0.08 ^*^ | 0.00 – 0.15 | **0.049** |
| Group | -0.36 ^***^ | -0.43 – -0.29 | **<0.001** |
| Reactivation Type * Valence | -0.01 | -0.04 – 0.02 | 0.480 |
| Reactivation Type * Group | 0.04 ^**^ | 0.01 – 0.07 | **0.010** |
| Valence * Group | -0.01 | -0.04 – 0.02 | 0.540 |
| Reactivation Type * Valence * Group | -0.02 | -0.05 – 0.01 | 0.228 |
| ** p<0.05   ** p<0.01   *** p<0.001* | | | |

*Table S15.* Results from linear mixed effects regression models predicting memory outcomes from emotional intensity ratings, reactivation type, and group. The model included random intercepts for subjects and videos, random slopes for *reactivation type* for both subjects and videos, and random slopes for *group* for videos.

| A) **Dependent Variable: Correct Details** | | | |
| --- | --- | --- | --- |
| *Predictors* | *Estimates* | *CI* | *p* |
| (Intercept) | -0.01 | -0.16 – 0.14 | 0.893 |
| Reactivation Type | -0.07 ^**^ | -0.12 – -0.02 | **0.007** |
| Intensity | -0.01 | -0.05 – 0.03 | 0.592 |
| Group | 0.16 ^*^ | 0.02 – 0.31 | **0.036** |
| Reactivation Type * Intensity | -0.03 | -0.08 – 0.02 | 0.248 |
| Reactivation Type * Group | -0.01 | -0.04 – 0.02 | 0.487 |
| Intensity * Group | -0.00 | -0.03 – 0.03 | 0.888 |
| Reactivation Type * Intensity * Group | -0.01 | -0.04 – 0.02 | 0.660 |
| **B) Dependent Variable: False Memories** | | | |
| *Predictors* | *Estimates* | *CI* | *p* |
| (Intercept) | 0.01 | -0.09 – 0.11 | 0.859 |
| Reactivation Type | -0.05 ^**^ | -0.08 – -0.02 | **0.001** |
| Intensity | -0.02 | -0.10 – 0.05 | 0.540 |
| Group | -0.36 ^***^ | -0.43 – -0.29 | **<0.001** |
| Reactivation Type * Intensity | 0.00 | -0.03 – 0.04 | 0.765 |
| Reactivation Type * Group | 0.04 ^**^ | 0.01 – 0.07 | **0.009** |
| Intensity * Group | -0.01 | -0.04 – 0.02 | 0.613 |
| Reactivation Type * Intensity * Group | 0.01 | -0.02 – 0.04 | 0.365 |
| ** p<0.05   ** p<0.01   *** p<0.001* | | | |

*Table S16.* Results from linear mixed effects regression models predicting memory outcomes from memorability ratings, reactivation type, and group. The model included random intercepts for subjects and videos, random slopes for *reactivation type* for both subjects and videos, and random slopes for *group* for videos.

| A) **Dependent Variable: Correct Details** | | | |
| --- | --- | --- | --- |
| *Predictors* | *Estimates* | *CI* | *p* |
| (Intercept) | -0.01 | -0.16 – 0.14 | 0.893 |
| Reactivation Type | -0.07 ^**^ | -0.12 – -0.02 | **0.008** |
| Memorability | -0.01 | -0.05 – 0.04 | 0.780 |
| Group | 0.16 ^*^ | 0.02 – 0.31 | **0.036** |
| Reactivation Type * Memorability | -0.01 | -0.06 – 0.04 | 0.764 |
| Reactivation Type * Group | -0.01 | -0.04 – 0.02 | 0.488 |
| Memorability * Group | 0.01 | -0.02 – 0.04 | 0.342 |
| Reactivation Type * Memorability * Group | -0.02 | -0.05 – 0.01 | 0.256 |
| **B) Dependent Variable: False Memories** | | | |
| *Predictors* | *Estimates* | *CI* | *p* |
| (Intercept) | 0.01 | -0.09 – 0.11 | 0.850 |
| Reactivation Type | -0.05 ^***^ | -0.08 – -0.02 | **0.001** |
| Memorability | -0.06 | -0.14 – 0.01 | 0.104 |
| Group | -0.36 ^***^ | -0.43 – -0.29 | **<0.001** |
| Reactivation Type * Memorability | 0.04 ^*^ | 0.01 – 0.07 | **0.016** |
| Reactivation Type * Group | 0.04 ^**^ | 0.01 – 0.07 | **0.009** |
| Memorability * Group | 0.00 | -0.03 – 0.03 | 0.772 |
| Reactivation Type * Memorability * Group | 0.01 | -0.02 – 0.04 | 0.427 |
| ** p<0.05   ** p<0.01   *** p<0.001* | | | |

**Basal Forebrain tSNR**

The basal forebrain lies within a ventral portion of the brain that is susceptible to signal dropout in fMRI, due to the close proximity to the nasal cavity. In order to ensure that our effects were not driven by noise, we calculated the average temporal signal-to-noise (tSNR) statistic for each participant. Use FSL math utilities, we generated a whole-brain voxel-wise tSNR map by dividing the mean by the standard deviation. We then masked each subject’s tSNR map to calculate average tSNR within the native-space basal forebrain masks.

We found that although there was considerable individual variability in basal forebrain tSNR (*M* = 50, *SD* = 17.5), there were no statistical outliers. Furthermore, we also tested the robustness of our results if the five subjects with the lowest tSNR scores (more than one standard deviation below the mean) were excluded from analysis (Table S17). The statistical significance of our findings remained unchanged when the lowest tSNR subjects were excluded. Overall, we concluded that our basal forebrain findings were not driven by the low tSNR subjects, and thus we did not exclude any subjects from the analyses reported in the main text.

*Table S17.* Analysis testing whether basal forebrain results held when the five subjects with the lowest tSNR in the basal forebrain (>1SD below the mean) were excluded from the analysis. Compare to Table S12B. Parameter estimates from a linear mixed effects regression model predicting subsequent false memories from *reactivation type* (Full vs. Interrupted), univariate *hippocampal activation* (HPC), post-video *change in hippocampal autocorrelation* (Autocor Change), univariate *basal forebrain activation* (BF), *hemisphere*, *axis*, and interaction terms. Model included random intercepts for subjects and videos; random slopes for reactivation type, univariate hippocampal activation, univariate basal forebrain activation, and autocor change for each subject; and random slopes for reactivation type for each video. Excluding the 5 subjects with the lowest (tSNR) in the basal forebrain did not change the key results (highlighted rows).

| **Dependent Variable: False Memories** | | | |
| --- | --- | --- | --- |
| *Predictors* | *Estimates* | *CI* | *p* |
| (Intercept) | 0.03 | -0.14 – 0.19 | 0.739 |
| Reactivation Type | -0.10 ^***^ | -0.14 – -0.06 | **<0.001** |
| BF | -0.01 | -0.06 – 0.05 | 0.831 |
| HPC | -0.04 ^*^ | -0.07 – -0.01 | **0.024** |
| Autocor Change | 0.00 | -0.03 – 0.04 | 0.876 |
| Hemisphere | 0.00 | -0.02 – 0.03 | 0.937 |
| Axis | 0.00 | -0.02 – 0.03 | 0.704 |
| Reactivation Type * BF | -0.02 | -0.04 – 0.01 | 0.290 |
| Autocor Change * Hemisphere | 0.01 | -0.02 – 0.03 | 0.686 |
| Autocor Change * Axis | -0.00 | -0.03 – 0.03 | 0.926 |
| Reactivation Type * HPC | -0.04 ^**^ | -0.07 – -0.02 | **0.002** |
| Reactivation Type * Autocor Change | 0.01 | -0.01 – 0.04 | 0.328 |
| Reactivation Type * Hemisphere | -0.00 | -0.03 – 0.02 | 0.964 |
| Reactivation Type * Axis | 0.00 | -0.02 – 0.03 | 0.769 |
| BF * HPC | -0.02 | -0.05 – 0.00 | 0.091 |
| BF * Autocor Change | 0.03 | -0.00 – 0.06 | 0.059 |
| BF * Hemisphere | -0.00 | -0.03 – 0.02 | 0.786 |
| BF * Axis | 0.00 | -0.02 – 0.03 | 0.738 |
| Reactivation Type * Autocor Change * Hemisphere | -0.02 | -0.04 – 0.01 | 0.251 |
| Reactivation Type * Autocor Change * Axis | -0.02 | -0.04 – 0.01 | 0.235 |
| BF * Autocor Change * Hemisphere | -0.01 | -0.03 – 0.02 | 0.690 |
| BF * Autocor Change * Axis | -0.01 | -0.04 – 0.02 | 0.566 |
| Reactivation Type * BF * HPC | -0.03 ^*^ | -0.05 – -0.00 | **0.028** |
| Reactivation Type * BF * Autocor Change | 0.04 ^**^ | 0.01 – 0.07 | **0.004** |
| Reactivation Type * BF * Hemisphere | -0.00 | -0.03 – 0.02 | 0.728 |
| Reactivation Type * BF * Axis | 0.01 | -0.02 – 0.03 | 0.589 |
| Reactivation Type * BF * Autocor Change * Hemisphere | -0.00 | -0.03 – 0.03 | 0.950 |
| Reactivation Type * BF * Autocor Change * Axis | 0.01 | -0.02 – 0.03 | 0.651 |
| ** p<0.05   ** p<0.01   *** p<0.001* | | | |

**Autocorrelation Control**

To determine the anatomical specificity of our autocorrelation findings, we tested two control regions: inferior lateral occipital cortex (LOC) and white matter. We predicted that autocorrelation in LOC would be sensitive to all video offsets because of the change in visual input, but *not* sensitive to prediction error. In contrast, physiological noise from white matter should not be sensitive to either video offsets or prediction errors. Autocorrelation in LOC significantly increased after videos (*t*(23) = 6.17, *p* < .001, Cohen’s *d* = 1.29, 95% CI [0.73, 1.83]), but did not differ by reactivation type (*t*(23) = -0.30, *p* = .766, *d* = -0.06, 95% CI [-0.47, 0.35]). Autocorrelation in white matter did not change post-offset (*t*(23) = 1.07, *p =* .294, *d* = 0.22, 95% CI [-0.19, 0.64]) and did not differ by reactivation type (*t*(23) = 0.82, *p =* .42, *d* = 0.17, 95% CI [-0.24, 0.58]). In summary, these control analyses indicated that our autocorrelation findings were not a brain-wide phenomenon.

**Video Duration Control**

Prolonged visual stimulation during video playback could influence the magnitude and duration of the BOLD response in the hippocampus (and elsewhere) after video offset. Because Full videos are, on average, longer in duration than Interrupted videos, it is important to rule out video duration as a confounding variable.

Our stimulus set included a varied range of video durations for both Full and Interrupted videos, making it possible to overcome this confound. We conducted a control analysis to equate video duration between the Full and Interrupted conditions. First, we took a subset of our data that omitted the longest Full videos (>= 38 seconds) and the shortest Interrupted videos (<= 27 seconds). Omitting these trials equated video duration for the Full and Interrupted conditions (Full mean = 31.6 s, Interrupted mean = 31.9 s), such that there was no significant difference in durations between conditions (β = -0.03, 95% CI [-0.06, 0.01], *t* = -1.47, *p* = .142). Note that although there was still a small, non-significant numerical difference in video durations between conditions, the direction of the difference was reversed, such that the average Interrupted video was slightly longer than the average Full video. These trial exclusions left between 17-29 viable trials per condition for each subject.

Using this subset of our data, we were able to reproduce all of our key findings, demonstrating that there was still a robust effect of reactivation type on the relationship among hippocampal activation, basal forebrain activation, and subsequent memory.

We conducted mixed effects linear regression to predict subsequent *false memories* from the variables *reactivation type* (Full vs. Interrupted), univariate *hippocampal activation,* post-video change in *hippocampal autocorrelation*, univariate *basal forebrain activation,* and all interactions of interest. We included covariates for *hemisphere* and *long-axis* position (for hippocampal ROIs) and random effects to account for variance by subject and video. The construction of the model was identical to the model reported in Table S12B. Notably, this model simultaneously reproduces all of our key findings, because it is an expanded version of the simpler models that were reported elsewhere in the Results. All parameter estimates are provided in Table S18.

We reproduced our univariate results with this subset of our data that controlled for video duration. There was a significant interaction between reactivation type and univariate hippocampal activation predicting subsequent false memories (β = -0.03, 95% CI [-0.06, -0.01], *t* = -2.35, *p* = .019). There was also a significant three-way interaction among reactivation type, univariate hippocampal activation, and basal forebrain activation predicting subsequent false memories (β = -0.04, 95% CI [-0.06, -0.01], *t* = -2.77, *p* = .006).

Additionally, we reproduced our autocorrelation findings. There was a significant interaction between reactivation type and post-video change in hippocampal autocorrelation predicting subsequent false memories (β = 0.05, 95% CI [0.02, 0.08], *t* = 3.47, *p* < .001). There was also a significant three-way interaction among reactivation type, post-video change in hippocampal autocorrelation, and basal forebrain activation predicting subsequent false memories (β = 0.03, 95% CI [0.01, 0.06], *t* = 2.28, *p* = .023).

Lastly, in a separate model we also tested whether video duration was associated with post-video change in autocorrelation, because this measure showed a difference between Full and Interrupted videos. Video duration was not significantly related to post-video autocorrelation (β = 0.01, 95% CI [-0.02, 0.04], *t* = 0.57, *p* = .574). Taken together, these control analyses provide compelling evidence that our results cannot be explained by the duration of video stimulation alone.

*Table S18*. Analysis controlling for video duration reproduces all key findings. Parameter estimates from linear mixed effects regression models relating hippocampal autocorrelation to subsequent false memories, using a subset of the data selected to equate the average video duration for Full and Interrupted videos. These results correspond to Table S12B, but reflect a subset of our data in which the average video durations for Full and Interrupted videos were equated. The model predicted subsequent false memories from the variables reactivation type (Full vs. Interrupted), univariate hippocampal activation (HPC), post-video change in hippocampal autocorrelation (Autocor Change), univariate basal forebrain activation (BF), hemisphere, axis, and relevant interaction terms. The model also included random intercepts for subjects and videos; random slopes for reactivation type, univariate hippocampal activation, autocorrelation change, and basal forebrain activation for each subject; and random slopes for reactivation type for each video. Boldface indicates statistically significant parameters. Highlighted rows indicate key findings that were reproduced in this subset of the data.

| **Dependent Variable: False Memories** | | | |
| --- | --- | --- | --- |
| *Predictors* | *Estimates* | *CI* | *p* |
| (Intercept) | 0.05 | -0.11 – 0.21 | 0.575 |
| Reactivation Type | -0.11 ^**^ | -0.17 – -0.05 | **0.002** |
| BF | -0.03 | -0.08 – 0.03 | 0.368 |
| HPC | -0.01 | -0.05 – 0.03 | 0.594 |
| Autocor Change | -0.03 | -0.07 – 0.01 | 0.143 |
| Hemisphere | -0.00 | -0.03 – 0.03 | 0.958 |
| Axis | 0.00 | -0.02 – 0.03 | 0.813 |
| Reactivation Type * BF | 0.01 | -0.02 – 0.04 | 0.628 |
| Autocor Change * Hemisphere | 0.02 | -0.01 – 0.05 | 0.184 |
| Autocor Change * Axis | -0.00 | -0.03 – 0.03 | 0.838 |
| Reactivation Type * HPC | -0.03 ^*^ | -0.06 – -0.01 | **0.019** |
| Reactivation Type * Autocor Change | 0.05 ^***^ | 0.02 – 0.08 | **0.001** |
| Reactivation Type * Hemisphere | 0.00 | -0.03 – 0.03 | 0.999 |
| Reactivation Type * Axis | 0.00 | -0.02 – 0.03 | 0.759 |
| BF * HPC | -0.01 | -0.03 – 0.02 | 0.708 |
| BF * Autocor Change | 0.02 | -0.01 – 0.04 | 0.312 |
| BF * Hemisphere | -0.00 | -0.03 – 0.03 | 0.998 |
| BF * Axis | 0.00 | -0.03 – 0.03 | 0.880 |
| Reactivation Type * Autocor Change * Hemisphere | -0.03 | -0.06 – 0.00 | 0.070 |
| Reactivation Type * Autocor Change * Axis | -0.02 | -0.04 – 0.01 | 0.298 |
| BF * Autocor Change * Hemisphere | -0.01 | -0.03 – 0.02 | 0.634 |
| BF * Autocor Change * Axis | -0.02 | -0.05 – 0.01 | 0.112 |
| Reactivation Type * HPC * BF | -0.04 ^**^ | -0.06 – -0.01 | **0.006** |
| Reactivation Type * Autocor Change * BF | 0.03 ^*^ | 0.00 – 0.06 | **0.023** |
| Reactivation Type * BF * Hemisphere | -0.00 | -0.03 – 0.03 | 0.884 |
| Reactivation Type * BF * Axis | 0.01 | -0.02 – 0.03 | 0.614 |
| Reactivation Type * Autocor Change * BF * Hemisphere | -0.01 | -0.04 – 0.02 | 0.532 |
| Reactivation Type * Autocor Change * BF * Axis | 0.01 | -0.02 – 0.04 | 0.389 |
| ** p<0.05   ** p<0.01   *** p<0.001* | | | |
